# PROFILING OF Y1 CELLS TREATED WITH FGF-2 REVEALS PARALLELS WITH ONCOGENE-INDUCED SENESCENCE

**DOI:** 10.1101/2020.08.17.247023

**Authors:** Peder J. Lund, Mariana Lopes, Simone Sidoli, Mariel Coradin, Francisca Nathália de Luna Vitorino, Julia Pinheiro Chagas da Cunha, Benjamin Aaron Garcia

## Abstract

Paradoxically, oncogenes that drive cell cycle progression may also trigger pathways leading to senescence, thereby inhibiting the growth of tumorigenic cells. Along these lines, Y1 cells, which carry an amplification of Ras, become senescent after treatment with the mitogen FGF-2. To understand how FGF-2 promotes senescence, we profiled the epigenome, transcriptome, proteome, and phospho-proteome of Y1 cells stimulated with FGF-2. FGF-2 caused delayed acetylation of histone H4 and higher levels of H3K27me3. Sequencing analysis revealed decreased expression of cell cycle-related genes with concomitant loss of H3K27ac. In contrast, FGF-2 promoted the expression of p21, various cytokines, and MAPK-related genes. Nuclear envelope proteins, particularly lamin B1, displayed increased phosphorylation in response to FGF-2. Proteome analysis suggested alterations in cellular metabolism, as evident by modulated expression of enzymes involved in purine biosynthesis, tRNA aminoacylation, and the TCA cycle. Altogether, the response of Y1 cells to FGF-2 is consistent with oncogene-induced senescence. We propose that Y1 cells enter senescence due to deficient cyclin expression and high levels of p21, which may stem from DNA damage or TGFb signaling.

## INTRODUCTION

The term “replicative senescence” was proposed by Hayflick and Moorhead in 1961 (Hayflick & Moorhead, 1961), referring to the inability of cultured cells to continue dividing indefinitely. Subsequent studies attributed this phenomenon to the shortening of telomeres (Bodnar *et al*, 1998). Unlike replicative senescence, oncogene-induced senescence (OIS) does not involve telomere shortening but rather the activation of oncogenes, such as Ras, Raf, E2F1, and MEK (Dimauro & David, 2010). Since oncogenes have established roles in promoting cell proliferation and tumorigenesis, OIS may seem counterintuitive at first but likely constitutes a cellular defense mechanism that guards against the development of cancer. Thus, investigating the pathways that underlie OIS, and how tumor cells circumvent those pathways, is important for understanding the process of tumorigenesis (Campisi, 2001).

In contrast to its typical activity as a mitogen, fibroblast growth factor 2 (FGF-2) has been observed to induce senescence in a number of cell lines. For instance, in the breast cancer cell line MCF-7, even though FGF-2 delivers mitogenic signals through ERK, resulting in increased levels of cyclin E, cyclin D1, and cyclin-dependent kinase 4, cells ultimately fail to proliferate due to increased levels of the cell cycle inhibitor p21 (Fenig *et al*, 1997). Similarly, after treatment with FGF-2, the neuroepithelioma cell line SK-N-MC arrests in G2 phase because Cdk1 (Cdc2) retains inhibitory phosphorylation at Tyr15, thereby preventing full activation of the cyclin B-Cdk1 complex (Smits *et al*, 2000). FGF-2 also causes senescence in the murine adrenocortical carcinoma cell line Y1 (Costa *et al*, 2008), which carries an amplification of the K-Ras protooncogene (Schwab *et al*, 1983). Treatment with serum causes Y1 cells to proliferate, but when serum is combined with FGF-2, Y1 cells transiently arrest in the G1/S transition and then experience an irreversible blockade at the G2/M transition that is consistent with senescence based on expression SA-beta-galactosidase (Costa *et al*, 2008). This phenotype can also be recapitulated in a mouse model, in which intradermal injections of FGF-2 suppress the formation of tumors that arise from inoculated Y1 cells (Costa *et al*, 2008). Notably, studies have demonstrated that the anti-proliferative effect of FGF-2 is dependent on Ras activity. Disruption of K-Ras through genome editing protects Y1 cells from senescence (Dias *et al*, 2019), as does expression of a dominant-negative Ras mutant (Costa *et al*, 2008). Conversely, 3T3 cells become susceptible to senescence after expression of the H-Ras^V12^ oncogene (Salotti *et al*, 2013). Through FGFR, FGF-2 triggers the activation of four major signaling pathways, including Ras-Raf-MAPK, PI3K-Akt, STATs, and PLC/PKC (Turner & Grose, 2010). Based on experiments with various inhibitors, the MEK, PI3K, and PKC pathways appear dispensable for the senescence phenotype, which was instead found to be dependent on RhoA and Src activity (Salotti *et al*, 2013). However, a more recent study, using a more potent inhibitor, provided some evidence of MEK involvement (Dias *et al*, 2019).

Given its dependence on Ras, the ability of FGF-2 to promote senescence in Y1 cells appears similar to OIS. As outlined by Dimauro and David (Dimauro & David, 2010), Ras-driven OIS consists of three primary features, which include: (i) formation of heterochromatin and transcriptional repression of E2F target genes by p16^Ink4a^ and pRb; (ii) activation of the DNA damage response from replicative stress; and (iii) expression of a senescence-associated secretory phenotype (SASP). Indeed, FGF-2 has been reported to cause an OIS-like phenotype in chondrocytes (Krejci *et al*, 2010). In addition to FGF-2, other growth factors are also capable of driving senescence. For instance, EGF promotes senescence in melanocytes expressing an oncogenic version of EGFR (Leikam *et al*, 2008). While this EGF-dependent senescence is proposed to result from DNA damage caused by ROS, a detailed mechanism explaining why Y1 cells senesce in response to FGF-2 is lacking. Thus, we aimed to uncover the molecular pathways that contribute to the senescence phenotype by applying genomics, transcriptomics, and proteomics to the study of Y1 cells treated with FGF-2. We find that FGF-2 leads to increased levels of H3K27me3, repression of cyclins and other E2F target genes, and expression of a number of soluble factors and cytokines, all of which are consistent with OIS. Altogether, our data has established the foundation of a working model in which FGF-2 triggers senescence in Y1 cells because of an inability to progress through the cell cycle from deficient cyclin expression and high levels of p21, which may be a downstream consequence of DNA damage or TGFb signaling.

## RESULTS

### FGF-2 delays acetylation on histone H4 and promotes repressive methylation on histone H3

Biological processes involving chromatin, including gene expression, are regulated in part by the deposition and removal of post-translational modifications (PTMs) on histone proteins. Thus, based on previous work demonstrating that senescence is accompanied by changes in the epigenetic landscape (Narita *et al*, 2003; Dimauro & David, 2010; Funayama *et al*, 2006; Parry & Narita, 2016), we hypothesized that cellular senescence induced by FGF-2 may also be accompanied by global alterations in histone modification levels, such as a reduction in marks related to transcriptional activation and an increase in marks related to transcriptional repression. To test this hypothesis, we treated G0-arrested Y1 cells with FBS alone or with FGF-2 and then analyzed histone PTMs at various timepoints after cell cycle progression by bottom-up mass spectrometry, which has emerged as an ideal analysis platform due to its high sensitivity, high throughput, and ability to resolve the many different types and sites of histone modifications. The timepoints chosen for analysis roughly correspond to the G1 (0.5-8 hrs), S (12-16 hrs), and G2/M (24 hrs) phases of the cell cycle based on findings with the derivative cell line Y1D1 (Dias *et al*, 2019). The full assortment of modifications and their relative abundances on two important regions of histone H3 (aa 27-40) and histone H4 (aa 4-17) are presented in detail in Supplemental Figs. 1 and 2. PTMs from other regions of H3 and H4 were also detected but were not remarkably affected by FGF-2 treatment. As depicted in Fig. 1A, Y1 cells in both conditions showed a gradual increase in the levels of di-, tri-, and tetra-acetylation on the peptide GKGGKGLGKGGAKR, which forms part of the N-terminal tail of histone H4 (amino acids 4-17) and contains four potential acetylation sites (H4K5, H4K8, H4K12, H4K16). Acetylation at this region is typically associated with chromatin accessibility, transcriptional activation, and cell cycle progression (Zhao & Garcia, 2015). However, cells treated with FGF-2 showed a delayed accumulation of histone H4 acetylation, most notably for the di-acetylated state, at times representing the transition between G1 and S (8-12 hrs), consistent with the delayed onset of S phase after FGF-2 treatment (Dias *et al*, 2019). In addition to changes in acetylation on histone H4, we also observed higher levels of tri-methylation at histone H3, lysine 27 (H3K27me3) in Y1 cells treated with FGF-2 for 24 hrs (Fig. 1B). In contrast to H4 acetylation, H3K27me3 is generally associated with inaccessible heterochromatin and transcriptional repression. Altogether, these results suggest that FGF-2 is associated with a delay in the accumulation of an activating mark on histone H4 and higher levels of a repressive mark on histone H3.

**Figure 1.**
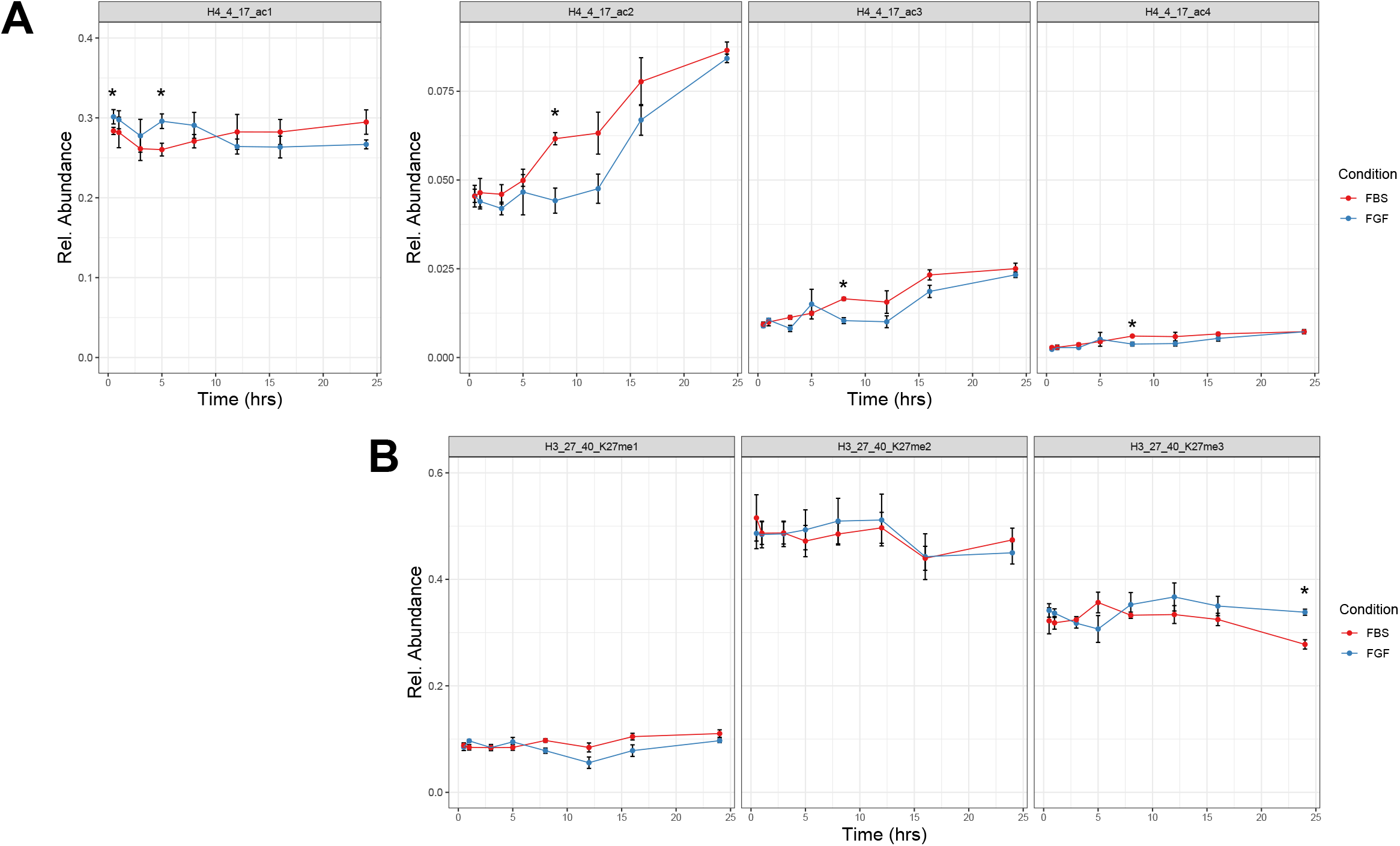
FGF-2 delays acetylation on histone H4 and promotes repressive methylation on histone H3. A) Histones were extracted from Y1 cells treated with FBS or FBS + FGF-2 for various amounts of time (0.5, 1, 3, 5, 8, 12, 16, 24 hrs). The relative abundance of the N-terminal histone H4 peptide (amino acids 4-17) carrying one, two, three, or four acetyl groups in total, irrespective of the precise position (K5, K8, K12, or K16), is plotted over time for the FBS and FGF conditions (mean ± s.e.m, n = 4). * p < 0.05 by unpaired t-test. B) The relative abundance of the H3 peptide (amino acids 27-40) carrying one, two, or three methyl groups at H3K27, irrespective of proximal methylation events at H3K36, is plotted over time for the FBS and FGF conditions (mean ± s.e.m, n = 4). * p < 0.05 by unpaired t-test.

### Altered expression of MAPK-related genes and cyclins in response to FGF-2 correlates with H3K27ac levels

Information concerning the genomic context of histone modifications is crucial for understanding how epigenetic alterations contribute to phenotypic outcomes, such as senescence. Furthermore, subtle but meaningful changes occurring at small subsets of genes may escape detection when analyzing histone modifications *en masse*. Thus, to complement our global histone PTM results, we next applied next-generation sequencing approaches to investigate how FGF-2 influences histone modifications at specific genes and the transcription of those genes. Given its established importance as a marker of enhancers and active promoters (Creyghton *et al*, 2010), we selected acetylation at H3K27 (H3K27ac) as the modification to profile by ChIP-seq, even though statistically significant changes were not apparent at the global level as assessed by mass spectrometry aside from the 8 hr timepoint, at which point the FGF condition had lower levels compared to the FBS condition (Supplemental Fig. 1). Because FGF-2 causes more than 70% of Y1 cells to enter senescence after 12 hrs (Costa *et al*, 2008), we opted for the earlier timepoint of 5 hrs to study the epigenetic events preceding cell cycle arrest and senescence.

Principal component analysis (PCA), whether considering all 40,783 consensus peaks or 3,426 differentially abundant peaks (Table 1), yielded strong separation across conditions but consistency within each condition (Supplemental Fig. 3A and 3B). Although the majority of peaks showed no quantitative differences between conditions, we observed 2,924 peaks with increased intensity in response to FGF-2 treatment and 502 peaks with decreased intensity (Supplemental Fig. 3C). Since global levels of H3K27ac show a downward trend in the FGF condition as assayed by mass spectrometry (Supplemental Fig. 1), a larger number of peaks showing increased versus decreased intensity is somewhat surprising, but it is possible that the downregulated peaks contribute a larger portion of the global H3K27ac signal due to differences in peak length or magnitude. When assigning peaks to genomic features, we noticed that proximal promoter regions (+/- 1 kb of TSS) had a higher representation among the peaks with decreased intensity after FGF-2 treatment compared to those with increased intensity (Supplemental Fig. 3D). In agreement with this trend, a more detailed investigation of transcription start sites (TSS) revealed substantially reduced H3K27ac signal intensity in the FGF-2 condition versus the FBS control (Fig. 2A), which indicates that FGF-2 either interferes with FBS-induced upregulation of H3K27ac near TSSs or promotes the removal of pre-existing H3K27ac marks.

**Figure 2.**
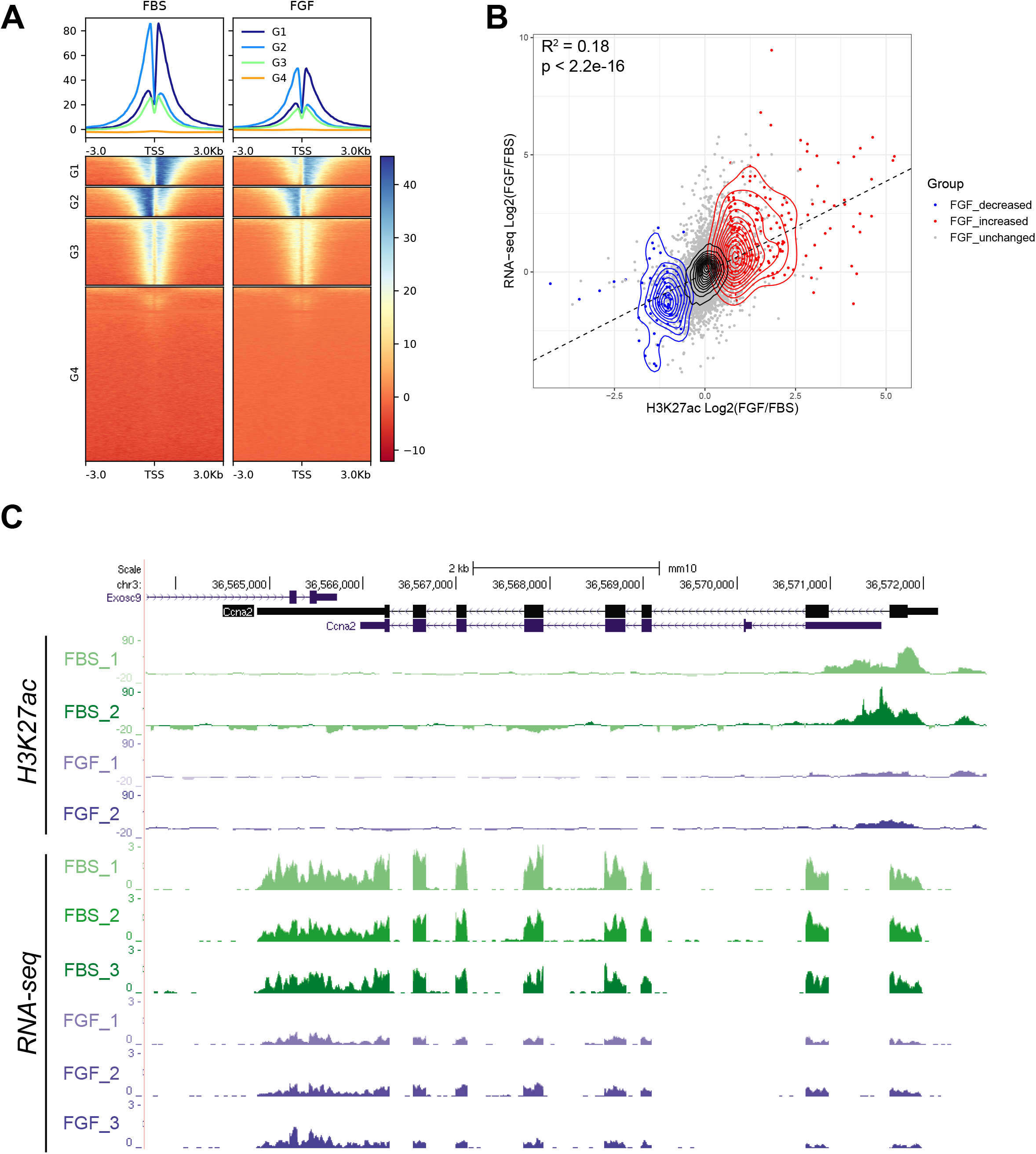
Altered expression of MAPK-related genes and cyclins in response to FGF-2 correlates with H3K27ac levels. A) The signal intensity of H3K27ac, normalized to input, is plotted over the regions surrounding unique TSS positions for the FBS and FGF conditions after 5 hrs of treatment. Replicates were merged for plotting. TSS regions are clustered into 4 groups (G1-4) by k-means based on signal intensity. B) The change in gene expression between the FGF and FBS conditions at 5 hrs, as assessed by RNA-seq, is plotted against the corresponding change in H3K27ac intensity. Each point represents a single gene (9,248 in total) and peaks were required to be within 10 kb of the TSS. For genes with multiple peaks, the peak closest to the TSS (and then the most intense peak in the case of ties) was considered. Contour plots are based only on changes in H3K27ac (q < 0.05 and log2(FGF/FBS) > 0 or < 0). Increases are shown in red and decreases are shown in blue. A regression line with the equation y = 0.785x - 0.055 (Pearson correlation R^2 = 0.18, p < 2.2e-16) is plotted. C) The *Ccna2* gene (cyclin-A2) is pictured along with tracks from the RNA-seq and H3K27ac ChIP-seq analyses. Replicates from the FBS and FGF conditions are shown in green and blue, respectively.

To assess whether changes in H3K27ac correlate with changes in transcription of proximal genes, we also performed RNA-seq in parallel. Consistent with prevailing dogma that H3K27ac associates with active genes, we observed a positive correlation between changes in H3K27ac status and changes in transcription levels (Fig. 2B and Table 2). We then carried out pathway enrichment analysis on the genes showing differentially abundant H3K27ac peaks as well as the genes showing concordant trends in the levels of both H3K27ac and RNA. Considering ChIP-seq data alone, the genes with decreased levels of H3K27ac after FGF-2 treatment (415 genes with log2[FGF/FBS] < -1 at FDR < 0.05) demonstrated enrichment for functions related to kinase activity and cell cycle regulation, among others (Supplemental Fig. 4A). Notable genes contained within these pathways included cyclins, regulatory subunits of protein kinase A, Raf1, and proteins involved in TGFb and Wnt signaling. On the other hand, genes with increased levels of H3K27ac after FGF-2 treatment (1421 genes with log2[FGF/FBS] > 1 at FDR < 0.05) showed an overwhelming enrichment for functions related to Ras and GTPase signaling and included a prominent number of guanine exchange factors and Rab family members (Supplemental Fig. 4B). In a second iteration of pathway analyses, we refined our focus to the subset of genes with matching trends in both H3K27ac and RNA levels. This revealed a functional enrichment for the MAPK cascade among the genes with increases in H3K27ac and RNA after FGF-2 treatment (157 genes with H3K27ac log2[FGF/FBS] > 0 at FDR < 0.05 and RNA log2[FGF/FBS] > 0 at p-adj < 0.05), and an enrichment for cell cycle regulation among the genes with decreases in H3K27ac and RNA (42 genes with H3K27ac log2[FGF/FBS] < 0 at FDR < 0.05 and RNA log2[FGF/FBS] < 0 at p-adj < 0.05) (Supplemental Fig. 5A and 5B). The former category featured several dual-specificity phosphatases (DUSPs) along with *Map3k11, Tgfb3, Fgf2*, and *Spry2.* Cyclins and genes related to spindle and kinetochore formation were noteworthy members of the latter category. Since FGF-2 treatment causes a delay and an eventual arrest in the cell cycle for Y1 cells, we were particularly interested in the cyclin genes, namely cyclin A2 (*Ccna2*) and cyclin B2 (*Ccnb2*), that exhibited differential expression between the FGF-2 and FBS conditions. As shown in Fig. 2C for *Ccna2* and Supplemental Fig. 5C for *Ccnb2*, the H3K27ac peaks observed near the 5’ end of the genes in the FBS condition are missing in the FGF-2 condition, concomitant with a reduction in the density of RNA-seq reads across the genes. Based on an integrated analysis of H3K27ac ChIP-seq and RNA-seq data sets from Y1 cells treated with FGF-2 for 5 hrs, we conclude that FGF-2 influences the levels of H3K27ac near distinct subsets of genes, which correlates with changes in expression of those genes. While genes related MAPK signaling gain H3K27ac and experience increased expression after FGF-2 treatment, genes related to cell cycle control lose or fail to gain H3K27ac and experience decreased expression.

### FGF-2 initiates a gene expression program indicative of an overactive MAPK cascade and a failure to prepare for cell division

While H3K27ac possesses clear biological importance, it represents only one of the many different types and sites of histone PTMs. As evident from Fig. 2B, significant changes in gene expression are possible without apparent changes in H3K27ac, perhaps as a result of other epigenetic modifications or the expression of specific transcription factors. Additionally, not every gene from our RNA-seq dataset matched to an annotated H3K27ac peak (of 11952 genes with valid adjusted p-values for RNA-seq, 9248 [77.4%] had H3K27ac peaks mapped within 10 kb of the TSS while 2204 [19.2%] of the 11452 genes with H3K27ac peaks mapping within 10 kb of the TSS had no or an inadequate number of RNA-seq reads to obtain a valid adjusted p-value). Therefore, we proceeded with a separate analysis of the RNA-seq data to gain a more complete understanding of how FGF-2 affects the transcriptome of Y1 cells compared to FBS alone (Table 3). As before, PCA easily distinguished between the two experimental conditions (Supplemental Fig. 6A). Heatmaps showing the genes with the most drastic fold-changes (adj. p < 0.05) are presented in Supplemental Fig. 6B. We performed gene set enrichment analysis (GSEA) and noted the emergence of several pathways bearing potential relevance to the senescence phenotype. FGF-2 treatment induced a transcriptional signature characterized in part by pathways involving cytokines, growth factors, and inactivation of MAPK signaling, among others (Fig. 3A). Indeed, Y1 cells showed increased expression of a number of growth factors and cytokines in response to FGF-2, including *Il11, Tgfb1, Pdgfb, Csf2*, and *Fgf2* itself (Fig. 3B, Supplemental Fig. 6C, Table 3). As mentioned in the previous section, numerous DUSPs were also upregulated. Supporting our earlier observation that FGF-2 suppresses the activation of cell cycle genes, GSEA indicated that Y1 cells in the FBS control condition were likely beginning to prepare for chromosome condensation and kinetochore formation based on increased expression of kinesins and centromere-related genes (Fig. 3C). Additionally, many other genes related to the cell cycle, and known to be regulated by the E2F family of transcription factors, were enriched in the FBS control condition, such as *Brca1, Ccnb2, Plk1*, and *Cenpe* (Fig. 3D). Notably, levels of *Cdkn1a*, which encodes the cell cycle inhibitor p21, were significantly lower in the FBS control condition (Supplemental Fig. 6C). Overall, we conclude that after 5 hrs of FGF-2 treatment, Y1 cells display a transcriptional signature consistent with the production of numerous growth factors and a disrupted ability to prepare for mitosis.

**Figure 3.**
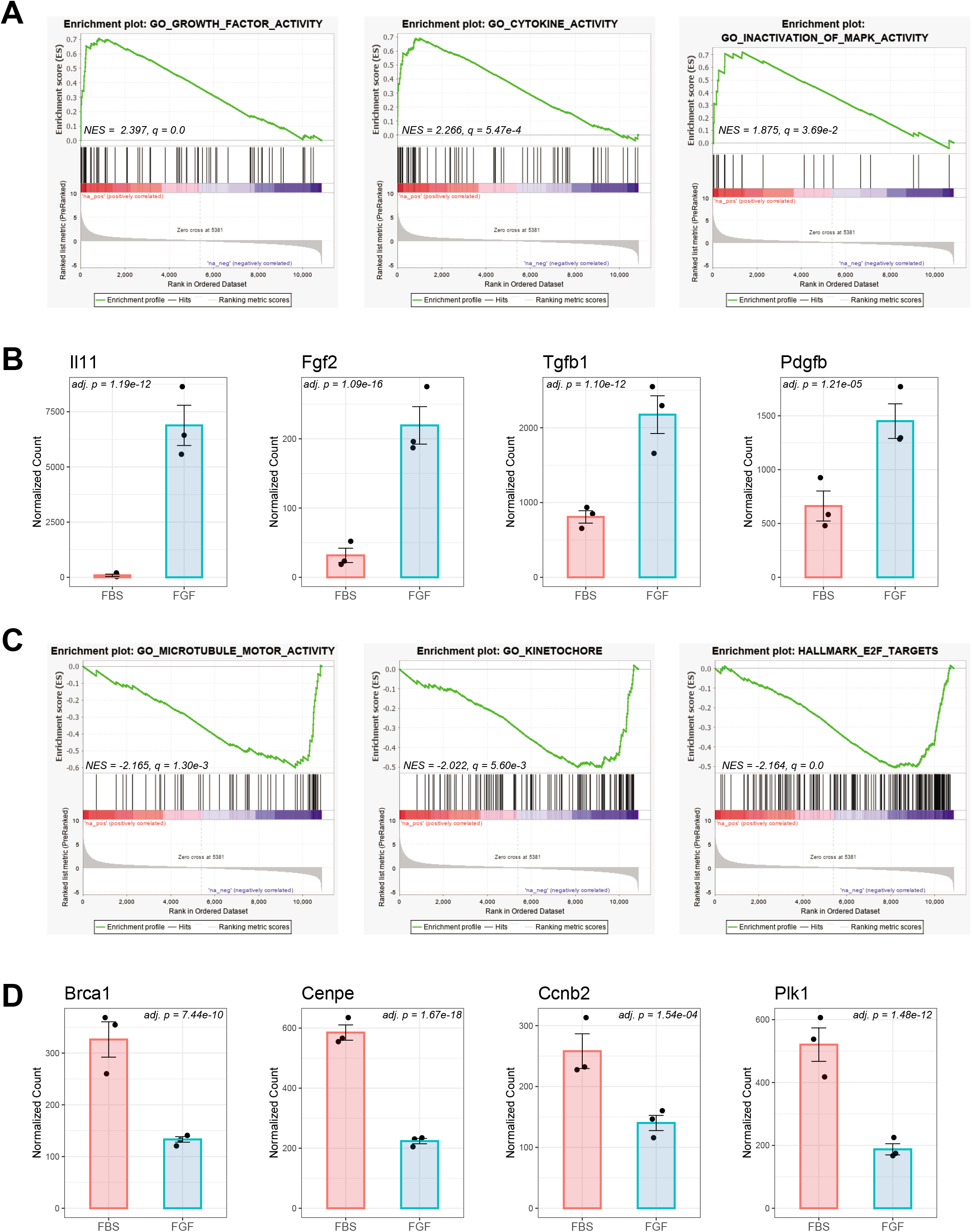
FGF-2 initiates a gene expression program indicative of an overactive MAPK cascade and a failure to prepare for cell division. A) RNA-seq data from Y1 cells treated with FBS or FBS + FGF for 5 hrs was analyzed by GSEA based on a ranked list of genes ordered by log2(FGF/FBS). Enrichment plots of GO terms with notable representation in the FGF condition are shown. Normalized enrichment score (NES) and FDR q-value (q) are shown from the GSEA. B) Normalized counts of several growth factors and cytokines are plotted for the FBS and FGF conditions. Bar charts display the group mean ± s.e.m. with individual replicates as points. Adjusted p-values are shown from the DESeq2 analysis. C) Enrichment plots of GO terms with notable representation in the FBS condition are shown. Normalized enrichment score (NES) and FDR q-value (q) are shown from the GSEA. D) Normalized counts of several cell cycle-related genes are plotted for the FBS and FGF conditions. Bar charts display the group mean ± s.e.m. with individual replicates as points. Adjusted p-values are shown from the DESeq2 analysis.

### FGF-2 leads to hyperphosphorylation of lamin B1 and modulates the expression of enzymes related to tRNA aminoacylation, purine biosynthesis, and oxidative respiration

To examine the upstream signaling events that precede the establishment of transcriptional and epigenetic programs that contribute to senescence, we next profiled protein phosphorylation patterns using mass spectrometry. Y1 cells were serum-starved and then treated with FBS alone or FBS with FGF-2 for 5, 10, 30, or 60 mins, all of which represent timepoints that correspond to G1 phase (Dias *et al*, 2019). To facilitate their identification, phospho-peptides were enriched from trypsin digests using titanium dioxide beads prior to analysis by LC-MS/MS (Tables 4-7). Consistently detected phospho-peptides with significant differences in protein-normalized intensities between the FBS and FGF-2 conditions at any timepoint are presented in Fig. 4A (see also Table 5). Normalization of phospho-peptides to the levels of their respective proteins is necessary to control for any changes in protein expression that could contribute to a perceived change in phospho-peptide abundance. Since phospho-peptides may also originate from less abundant proteins that lack intensity measurements and only minimal changes in protein abundance are expected at these early timepoints (Bahrami & Drabløs, 2016; Vitorino *et al*, 2018), we performed a second analysis of phospho-peptide intensities without protein normalization (Supplemental Fig. 7A, Table 6). To account for phospho-peptides detected solely, or mostly, in one condition, which complicates statistical calculations from intensity values, we conducted a third analysis using chi square tests (Supplemental Fig. 7B, Table 7). As expected, TiO2 beads resulted in preferential enrichment of phospho-serine over phospho-tyrosine (Supplemental Fig. 7C), which suggests that our data set may have limited representation of membrane-proximal tyrosine phosphorylation events. Hierarchical clustering of the samples in Fig. 4A and Supplemental Fig. 7A allowed for distinction between the FBS and FGF conditions at the later timepoints (30-60 mins) and to some extent at 10 mins. Focusing first on the protein-normalized phospho-peptides from Fig. 4A, we observed that FGF-2 treatment for 30 and 60 mins leads to hyperphosphorylation of several proteins involved in the breakdown of the nuclear envelope (Supplemental Fig. 7D), particularly at Thr21 and Ser24 of lamin B1 (Fig. 4B). Located in the N-terminal head domain of lamin B1, these mitotic phosphorylation sites are thought to prevent lamin polymerization, thereby destabilizing the nuclear lamina (Machowska *et al*, 2019; Kuga *et al*, 2010; Fiume *et al*, 2008). Although lacking normalization to protein levels, additional phospho-peptides of interest appear in Supplemental Fig. 7A. For example, after FGF-2 treatment, we detected increased levels of a phosphopeptide from Spred1, which belongs to the MAPK cascade. Phospho-peptides from Rrp1b, Baz1b, and Ahctf1 were also more abundant after FGF treatment. Both Baz1b and Rrp1b have been implicated in DNA damage response pathways (Xiao *et al*, 2009; Paik *et al*, 2010) while Ahctf1 is known to be involved in processes related to the nuclear lamina (Doucet *et al*, 2010; Rasala *et al*, 2006; Gillespie *et al*, 2007; Gao *et al*, 2011). In contrast, we found that FGF-2 treatment leads to decreased levels of a phospho-peptide from Trip12, which is an E3 ligase important for regulating p19Arf (Chen *et al*, 2010) and histone ubiquitination after DNA damage (Gudjonsson *et al*, 2012).

**Figure 4.**
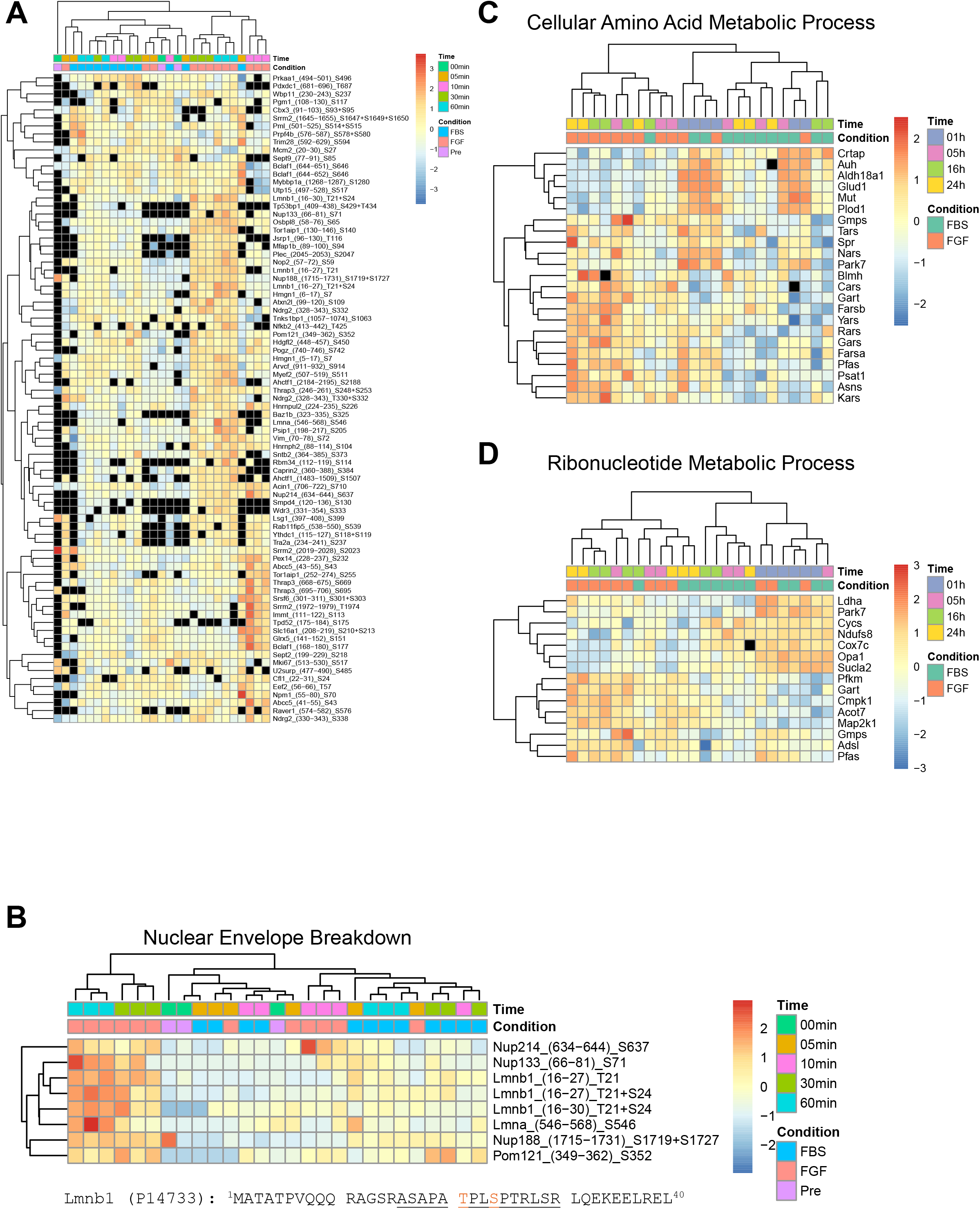
FGF-2 leads to hyperphosphorylation of lamin B1 and modulates the expression of enzymes related to tRNA aminoacylation, purine biosynthesis, and oxidative respiration. A) A heatmap of phospho-peptides with significant changes in at least one timepoint (p < 0.05, abs(Z-score of log2(FGF/FBS)) > 2, quantified in all replicates in both conditions for at least 1 timepoint) in intensities is presented. Phospho-peptide intensity is normalized to the intensity of the respective protein. Missing values, either from missing measurements of the phosphopeptide or its corresponding protein, are shown in black. Rows, which represent unique phospho-peptides, are named as the gene symbol with the corresponding amino acid positions and phosphorylation site. Cells were treated for 5, 10, 30, or 60 mins. “Pre” represents cells after serum starvation prior to treatment with FBS or FBS + FGF. B) A heatmap of phospho-peptides related to nuclear envelope breakdown and showing significant differences between the FBS and FGF conditions is displayed. Missing values were replaced with the row minimum to permit hierarchical clustering. Row names indicate the gene symbol, starting and ending positions of the peptide, and the phosphorylation sites. The first 40 amino acids of lamin B1 (Lmnb1) are shown below. The underlined portion corresponds to peptides detected by mass spectrometry with phosphorylation sites (Thr21 and Ser24) indicated in orange. C, D) Heatmaps of proteins related to the metabolism of amino acids (C) or ribonucleotides (D) based on GO analysis of are presented. Proteins for GO analysis (n = 248) and plotting were selected based on significant changes across the two conditions (p < 0.05 at either 16 h or 24 h timepoint and consistent sign in Z-scores at 5 h, 16 h, and 24 h timepoints).

In addition to profiling the phospho-proteome, we also employed mass spectrometry to monitor protein expression over time in Y1 cells treated with FBS alone or with FGF-2 (Tables 8 and 9). For this analysis, we focused on a wider range of timepoints to cover the G1 (1 and 5 hrs), S (16 hrs), and G2/M (24 hrs) phases of the cell cycle. Proteins showing significantly different abundances across conditions are depicted in Supplemental Fig. 8A. As evident by PCA, time accounts for more variation in the proteome than FGF-2 treatment (Supplemental Fig. 8B), which is most likely driven by changes in protein expression as cells transition out of serum starvation. However, the FGF-2 and FBS conditions achieved separation along PC2 at the later timepoints of 16 and 24 hrs, as the senescence phenotype becomes established. Pathway analysis of proteins showing differential abundance at these timepoints indicated possible regulation of multiple metabolic processes. Aminoacyl tRNA synthetases for threonine (Tars), asparagine (Nars), cysteine (Cars), tyrosine (Yars), phenylalanine (Farsa/b), arginine (Rars), glycine (Gars), and lysine (Kars) were upregulated after FGF-2 treatment (Fig. 4C), which could indicate a higher demand for protein translation, as previously proposed (Dias *et al*, 2019), or a compensatory response to inhibited translation. Two enzymes related to the synthesis of serine and asparagine, Psat1 and Asns, were also more abundant. Similarly, FGF-2 led to increased levels of enzymes related to purine biosynthesis, including Gart, Cmpk1, Gmps, Adsl, and Pfas (Fig. 4D). This finding may suggest an increased demand for purine nucleosides to support the production of cofactors for enzymatic reactions (e.g. ATP, GTP, NAD+) or precursors for nucleic acid synthesis. Finally, we noted that FGF-2 modulated the expression of several proteins involved in the TCA cycle and mitochondrial respiration (Supplemental Fig. 8C), such as Dlst, Sucla2, Aco2, Ndufb11, Ndufa13, and Ndufs8. Although these proteins appeared to be induced by serum starvation, and therefore their levels declined after returning cells to standard growth conditions, FGF-2 resulted in a more pronounced decrease compared to FBS alone. Notably, Ldha remained at higher levels in the FGF-2 condition. These results support the notion that mitochondrial metabolism declines as cells enter senescence. In summary, our analysis of the proteome and phospho-proteome by mass spectrometry suggests that FGF-2 treatment leads to hyperphosphorylation of lamin B1, upregulation of enzymes related to tRNA aminoacylation and purine biosynthesis, and downregulation of enzymes involved in mitochondrial metabolism.

## DISCUSSION

Fibroblast growth factor 2 (FGF-2) is often considered a mitogenic agent for different cell types and may even contribute to carcinogenesis (Turner & Grose, 2010). However, in murine adrenocortical carcinoma Y1 cells, which harbor an amplified and constitutively overexpressed K-Ras proto-oncogene (Schwab *et al*, 1983), FGF-2 causes delayed entry into and progression through S phase and an irreversible arrest at the G2/M transition, ultimately leading to cellular senescence (Dias *et al*, 2019; Costa *et al*, 2008). Here, we have employed a number of high-throughput technologies to investigate the potential mechanisms and pathways that underlie this atypical response. At the epigenetic level, Y1 cells treated with FGF-2 displayed higher levels of H3K27me3 and delayed acetylation of histone H4. Furthermore, the promoter regions of several genes related to the cell cycle displayed less H3K27ac, which correlated with lower transcription. In contrast, genes encoding growth factors, cytokines, and components of signaling cascades showed higher expression based on transcriptome analysis. At the protein level, FGF-2 led to the upregulation of enzymes involved in purine biosynthesis and tRNA aminoacylation and downregulation of enzymes related to mitochondrial metabolism at later timepoints (see Supplemental Discussion). Finally, representing one of the earliest divergences between conditions, FGF-2 resulted in the hyperphosphorylation of certain nuclear envelope proteins, particularly at the mitotic phosphorylation sites of lamin B1.

Although descriptive in nature, we believe our hypothesis-generating approach has enabled the synthesis of a more complete working model of the process by which FGF-2 causes senescence in Y1 cells (Fig. 5), which incorporates aspects of earlier studies (Costa *et al*, 2008; Salotti *et al*, 2013; Dias *et al*, 2019) and can be further refined by future, hypothesis-driven approaches. A key finding from our studies is that within 5 hours, Y1 cells treated with FGF-2 exhibited a transcriptional signature indicative of cell cycle arrest, highlighted by expression of p21 (*Cdkn1a*) and the repression of cyclins A2 and B2 (*Ccna2* and *Ccnb2*). Cyclins, together with cyclin-dependent kinases (Cdk), are critical regulators of cell cycle checkpoints in eukaryotic cells (Johnson & Walker, 1999). Ccna2 activates Cdk1 and is necessary for DNA replication during S phase and for M phase entry (Pagano *et al*, 1992). In contrast, p21 suppresses Cdk activity (Xiong *et al*, 1993; Georgakilas *et al*, 2017). By negatively regulating Cdks, p21 prevents phosphorylation of the retinoblastoma protein (Rb), resulting in continued sequestration of the E2F family transcription factors that are necessary for cell cycle progression (Bertoli *et al*, 2013; Fischer & Müller, 2017). More recently, inhibition of Cdk activity by p21 has also been shown to block phosphorylation of the Rb-related proteins Rbl1 (p107) and Rbl2 (p130). Similar to Rb, these proteins function as transcriptional repressors as part of the DREAM complex, which appears to control expression of an even larger subset of cell cycle genes than the E2F family (Engeland, 2018; Fischer *et al*, 2016a, 2016b). Closer inspection of our transcriptome data reveals strong evidence that cell cycle genes regulated by E2F and DREAM are repressed in cells treated with FGF-2 for 5 hrs (Fig. 3C and Supplemental Fig. 9B), consistent with the slower progression of Y1 cells through G1 and S phase as reported by others (Salotti *et al*, 2013; Dias *et al*, 2019). Accordingly, gene expression associated with the G1/S and G2/M transitions showed greater enrichment in the FBS control treatment (Supplemental Fig. 9B). Precisely what causes this early arrest remains unclear, but one possibility is the activation of p53 by DNA damage. Although an overall transcriptional signature indicative of p53 activity in the FGF condition was less apparent than for the genes regulated by E2F and DREAM (Fig. 3C and Supplemental Fig. 9), p21 is a direct transcriptional target of p53 (Jung *et al*, 2010; El-Deiry *et al*, 1993), and repression by the DREAM complex appears to depend on upstream expression of p21 mediated by p53 (Fischer *et al*, 2016a, 2016b). Furthermore, our examination of the transcriptome at a single timepoint (5 hrs) may have failed to capture an earlier peak in p53 activity, which is known to oscillate, potentially with a frequency and magnitude related to the type of DNA damage (Batchelor *et al*, 2011; Lahav *et al*, 2004), and to precede increases in p21 (Stewart-Ornstein & Lahav, 2016).

**Figure 5.**
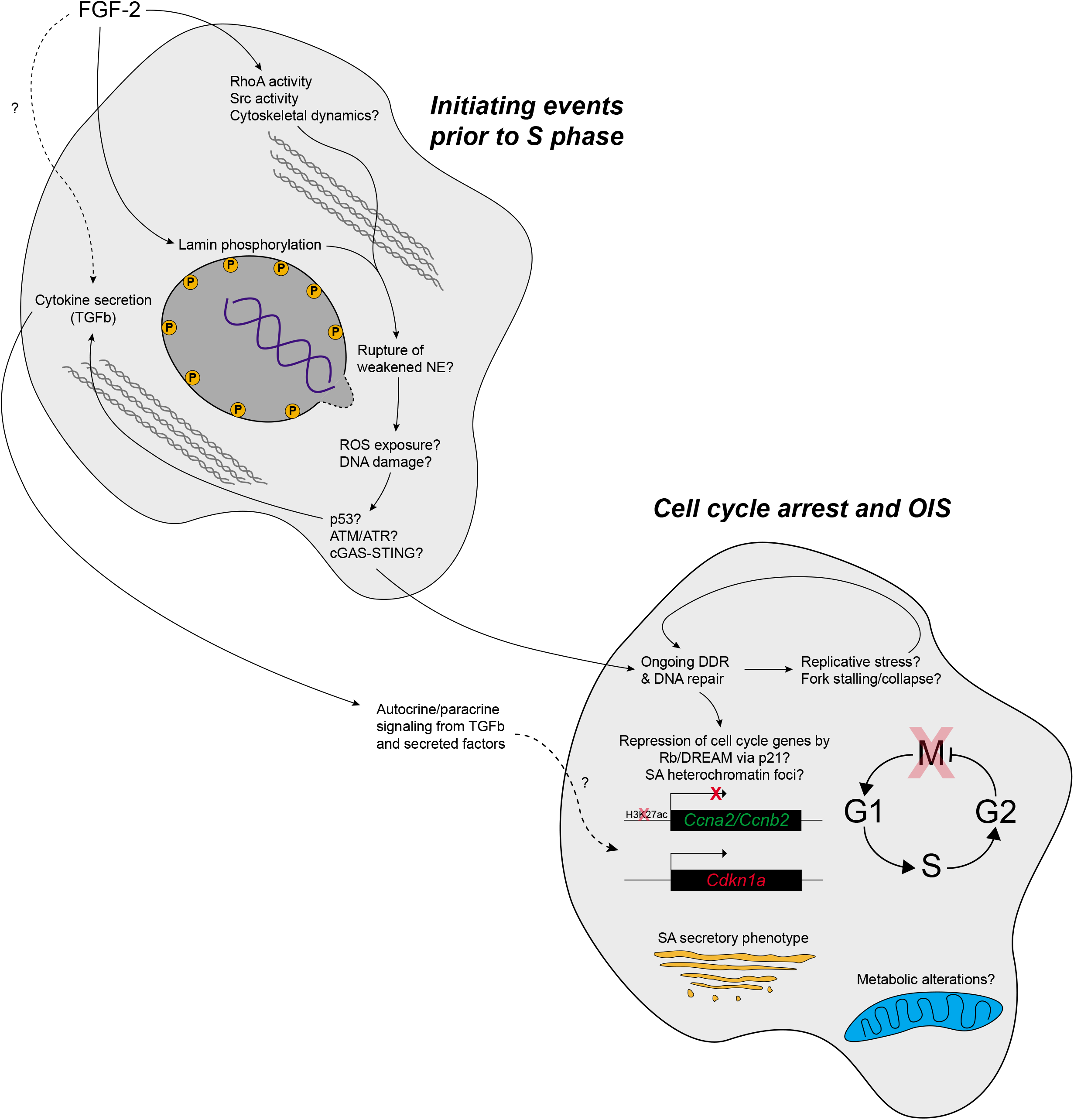
Proposed model of senescence in Y1 cells treated with FGF-2. In Y1 cells with amplified Ras, signaling from FGF-2 leads to increased cytoskeletal activity through RhoA (Salotti *et al*, 2013). Higher levels of phosphorylated lamin B1 in the nuclear envelope weaken its structural integrity, allowing transient rupture events from the mechanical stress of cytoskeletal remodeling. Exposed DNA is then prone to damage from cytosolic ROS, activating p53 and ATM/ATR. Cytosolic DNA may also activate cGAS-STING. Secreted factors, including TGFb, may be induced as a result of FGF-2 signaling or indirectly as a result of cell stress pathways. Downstream of the initiating events, an ongoing DNA damage response (DDR) and potentially autocrine/paracrine signaling from secreted factors, lead to expression of p21 and repression of cyclins, thereby enforcing cell cycle arrest. DDR is a hallmark of oncogene-induced senescence together with SASP, SAHF, and repression of E2F targets, possibly through the p53/p21/DREAM in addition to the more classical Rb pathway. Senescent cells may also feature metabolic deficiencies (reduced tRNA aminoacylation and ETC activity and increased purine biosynthesis), but it is unclear whether this is more of a cause or consequence of senescence.

Replicative stress during S phase may account for the DNA damage that is observed at later timepoints (Di Micco *et al*, 2006; Dimauro & David, 2010; Dias *et al*, 2019; Bartkova *et al*, 2006). However, since DNA synthesis is not evident until after 10-12 hrs of serum stimulation, at least as examined in Y1D1 cells that additionally have engineered expression of cyclin D (Dias *et al*, 2019), we suspect that an earlier event must exist to explain the DNA damage that we speculate to occur before S phase. From our phospho-proteome data, we noted phosphorylation sites on lamin B1 at Thr21 and Ser24 (Fig. 4B). These N-terminal phosphorylation sites in the head domain, along with additional phosphorylation sites in the C-terminal tail domain, are known to be targets of Cdk1 and potentially other kinases, such as PKC (Torvaldson *et al*, 2015; Machowska *et al*, 2019; Mall *et al*, 2012). During the normal events of mitosis, phosphorylation of these residues interferes with lamin polymerization, thereby destabilizing the nuclear lamina and allowing for nuclear envelope breakdown. In our system, N-terminal phosphorylation of lamin B1 occurs within 30-60 mins of FGF-2 treatment. We postulate that this early phosphorylation of lamin B1 weakens the integrity of the nuclear envelope. Recent work has demonstrated that mechanical forces, such as those experienced by cells undergoing migration, can result in transient rupturing of the nuclear envelope with subsequent damage to DNA (Hatch, 2018; Hatch & Hetzer, 2016; Denais *et al*, 2016; Lim *et al*, 2016; Raab *et al*, 2016). Interestingly, inhibition of RhoA, an important regulator of actin stress fibers and focal adhesions (Tojkander *et al*, 2012; Ridley & Hall, 1992), protects Y1 cells from FGF-2-induced senescence (Salotti *et al*, 2013; Costa *et al*, 2008). Given these previous observations, which included a mention of defective cell migration, we propose that FGF-2 leads to activation of RhoA and subsequent cytoskeletal remodeling, thereby generating mechanical forces that cause ongoing rupture of a nuclear envelope weakened by lamin phosphorylation. Upon nuclear rupture, the exposed DNA would then be susceptible to damage by cytosolic factors, such as nucleases and reactive oxygen species (ROS) (Lim *et al*, 2016). The latter is particularly relevant to the setting of FGF-2 and oncogenic Ras as multiple studies have already linked receptor signaling and Ras activation to the generation of ROS, and suppression of ROS can rescue cells from senescence (Irani *et al*, 1997; Sundaresan *et al*, 1996; Lee *et al*, 1999; Ogrunc *et al*, 2014; Leikam *et al*, 2008). We envision a scenario in which early oxidative damage introduces minor lesions in the form of oxidized bases or single-strand breaks, thereby delaying progression to S phase. Upon eventual entry into S phase, perhaps due to the high levels of mitogenic signaling, any unresolved damage could contribute to replicative stress and replication fork stalling (Zeman & Cimprich, 2014; Hills & Diffley, 2014). In turn, additional forms of DNA damage, including double-strand breaks, may then arise and lead to permanent arrest in G2/M. While another study provided evidence of DNA damage and ATR activation in Y1D1 cells at least as early as 24 hrs after FGF-2 treatment (Dias *et al*, 2019), which was attributed to replicative stress, a more precise examination of the kinetics and nature of DNA damage, as well as the involvement of specific DNA damage response and repair pathways and their contribution to the senescence phenotype, is necessary to either support or refute our proposed model.

The expression of p21 may also be linked to soluble factors, such as TGFb, that we found to be upregulated at the transcriptional level after FGF-2 stimulation. TGFb, which has been associated with DNA damage (Li *et al*, 2019) and senescence (Massagué *et al*, 2000; Seoane *et al*, 2004; Moustakas & Kardassis, 1998; Tian *et al*, 2011), activates the SMAD proteins to regulate gene expression affecting a wide array of cellular processes. In addition to those controlled by p53, the *Cdkn1a* gene also contains regulatory elements under control of SMADs through Sp1, and thus, TGFb is able to promote the expression of p21 (Li *et al*, 2019; Jung *et al*, 2010; Adnane *et al*, 1998; Lin *et al*, 2012). Strikingly, in at least one case, TGFb has even been found to induce senescence independent of p53 and the DNA damage response (Cipriano *et al*, 2011). Notably, after FGF-2 stimulation, Y1 cells downregulate expression of Smad6 (Supplemental Fig. 6B), which represents an inhibitory Smad (Jung *et al*, 2013). Thus, decreased levels of Smad6 may sensitize Y1 cells to TGFb signaling, resulting in the expression of p21 and cell cycle arrest. Pathway analysis of our transcriptome data further supports the relevance of TGFb signaling after FGF-2 treatment (Supplemental Fig. 9A). These findings invite a new perspective to interpreting an earlier study, which demonstrated that the PP1 inhibitor restores the ability of Y1 cells to grow after FGF-2 treatment (Salotti *et al*, 2013). Although this effect was attributed to inhibition of Src kinases downstream of Ras and RhoA, which could disrupt the cytoskeletal forces that we propose to be deleterious to the nuclear envelope (Tsuji *et al*, 2002; Yamana *et al*, 2006; Mitra *et al*, 2005), PP1 is also known to block TGFb signaling (Ungefroren *et al*, 2011; Maeda *et al*, 2006). Thus, the effect of PP1 may have occurred through Src, TGFb, or both.

Growth factor signaling is a ubiquitous and fundamental cellular process. Given that many growth factors share common signaling cascades downstream of their receptors, our findings on FGF-2 in the cellular context of Ras amplification may be relevant to other situations involving mitogenic stimulation of potentially oncogenic cells. Since FGF-2 signaling and the senescence phenotype both depend on Ras (Turner & Grose, 2010; Dias *et al*, 2019; Costa *et al*, 2008), we suspect that our model represents another instance of oncogene-induced senescence (OIS), a well-documented phenomenon that is thought to suppress the development of tumors from cells with activated oncogenes (Dimauro & David, 2010; Serrano *et al*, 1997). Previous work has highlighted three main features of Ras-driven OIS, namely (i) formation of senescence-associated heterochromatin foci (SAHF) and repression of E2F target genes via p16 and pRb, (ii) acquisition of a senescence-associated secretory phenotype (SASP), and (iii) activation of DNA damage response pathways due to replicative stress (Dimauro & David, 2010). Although we do not report direct morphological evidence of SAHF, our mass spectrometry analysis of histones demonstrated an increase in the repressive mark H3K27me3 after 24 hrs of FGF-2 stimulation, and we also found reduced levels of lamin B1 protein, as reported previously (Shimi *et al*, 2011) and which may affect SAHF formation (Sadaie *et al*, 2013). Consistent with the idea that SAHF formation coincides with repression of E2F target genes, our transcriptome analysis clearly indicated lower expression of many E2F targets and cell cycle-related genes after FGF-2 stimulation (Fig. 3C and Supplemental Fig. 9B), similar to a prior study of Ras-induced senescence (Mason *et al*, 2004). We also observed strong evidence of a SASP from our RNA-seq data. While the exact composition of SASP seems to vary according to cell type (Coppé *et al*, 2008), FGF-2 induced the expression of many previously reported SASP components, such as Plaur, Csf1, Fgf2, Il11, Pdgf, Tgfb, and PAI-1, as well as other cytokines, including Il6, Il24, Il33, and Ifnk (Fig. 3, Table 3, and additional notes in Supplemental Discussion). Peculiarly, the SASP appears to contain factors with opposite effects on cell growth, such as TGFb versus PDGF. A potential explanation for this dichotomy is that physiological levels of growth factor signaling drive the expression of positive regulators, possibly via the FOS, JUN or MYC families (Vitorino *et al*, 2018; Lepique *et al*, 2004), while oncogenic levels of growth factor signaling additionally induce the expression of negative regulators. The third and final feature of OIS is the activation of the DNA damage response, which has already been covered at length above. Based on the data presented here as well as in other publications, we believe that the senescence phenotype arising from the treatment of Y1 cells with FGF-2 is largely consistent with the general phenomenon of oncogene-induced senescence. From this perspective, the observation that a mitogen drives cells into a state of senescence instead of proliferation, though counterintuitive at first, is readily rationalized as the appropriate outcome due to the potentially oncogenic nature of Y1 cells, owing to probable DNA damage from the synergistic effect of FGF signaling in the background of K-Ras amplification. Continued study of how aberrant oncogenic stimulation pushes cells into a state of senescence will help to illuminate the ways in which tumor cells subvert these pathways, potentially sparking the development of novel strategies for the treatment of cancer.

## SUPPLEMENTAL DISCUSSION

In addition to TGFb, PAI-1 (encoded by *Serpine1*) is also upregulated by FGF-2-stimulated cells and likewise has been implicated in driving senescence (Serrano *et al*, 1997; Vaughan *et al*, 2017). Providing further evidence of early p53 activation in our model, previous studies have shown that p53 regulates the transcription of *Serpine1* (Kunz *et al*, 1995) and that PAI-1 is necessary for the induction of p53-dependent replicative senescence (Kortlever *et al*, 2006). Beyond classical DNA damage pathways, the cGAS-STING pathway, which is responsible for the sensing of cytosolic DNA, may also contribute to expression of SASP components. Cytosolic DNA has been previously reported in senescent cells (Glück *et al*, 2017; Yang *et al*, 2017; Dou *et al*, 2017; Li & Chen, 2018), and in a these studies, one of which included a model of senescence based on the HRasV12 oncogene, activation of the cGAS-STING pathway resulted in the expression of SASP components, such as the cytokines IL-6 and IFNb. Suggesting the presence of cytosolic DNA in our experimental model, Y1 cells expressed Il6 and Ifnk, another type I interferon like IFNb, in response to 5 hrs of FGF-2 stimulation (Table 3). We also observed the expression of chemokines, such as Ccl28, which, in an *in vivo* setting, may work in conjunction with these inflammatory cytokines to recruit and activate immune cells to eliminate potentially oncogenic cells (Dou *et al*, 2017).

Given the number of soluble factors expressed by Y1 cells in response to FGF-2, including Fgf2 itself, future studies should focus on which factors are important for the senescence phenotype and whether senescence is dependent on continued stimulation of growth factor receptors. Based on previous literature, we predict that ongoing stimulation is likely required. Dias and colleagues (Dias *et al*, 2019) noted sustained phosphorylation of ERK1/2 even after 24 hrs of FGF-2 stimulation, and inhibition of the MAPK pathway, which has been explored as a therapeutic target for tumors (Germann *et al*, 2017), appears to protect Y1 cells from senescence. In support of the importance of MAPK signaling, we observed higher expression of MAPK-related genes after FGF-2 treatment, including negative regulators, such as Spry2, Cnksr3, Dusp5, Dusp6, and Dusp7 (Supplemental Fig. 6 and Table 3). The induction of DUSP expression may act as a negative feedback mechanism in response to elevated MEK/MAPK activity, as seen in models of thyroid cancer with mutant BRAF (Buffet *et al*, 2017). Another study found that Dusp5 is a transcriptional target of p53, and as expected, DNA damage is able to drive p53-dependent expression of Dusp5 (Ueda *et al*, 2003). Furthermore, the overexpression of Dusp5 inhibited the growth of tumor cells. Thus, the expression of DUSPs may contribute to tumor suppression in the face of cellular stress, such as oncogenic signaling or DNA damage, and may be relevant to the mechanism by which FGF-2 induces senescence in Y1 cells.

As the senescence phenotype becomes more firmly established over time, our proteomic analysis at later timepoints (16-24 hrs) indicated possible alterations in cellular metabolism based on decreased expression of proteins related to mitochondrial respiration and increased expression of proteins related to purine biosynthesis and the charging of tRNA. From our data, we cannot distinguish whether this is a direct cause versus indirect consequence of senescence, but inhibition of oxidative phosphorylation has been reported to induce a senescent phenotype (Moiseeva *et al*, 2009). In either case, we speculate that the bioenergetic pathways in the senescing cells are deficient, which is consistent with previous reports of mitochondrial dysfunction in senescence (Moiseeva *et al*, 2009). Decreased production of NADH from the TCA pathway and less flux of NADH to complex I of the electron transport chain may lead to decreased production of ATP. The increased expression of enzymes involved in purine biosynthesis may represent an attempt to compensate by increasing the supply of the nucleotide precursors that give rise to ATP and other important cofactors, such as GTP and NAD. Another possible explanation is that unlike the control cells, which have completed cell division by 24 hrs (Dias *et al*, 2019), a subset of the senescing cells remain arrested in S phase and accordingly have upregulated the machinery to support DNA synthesis. A third possibility is that cells experience an increased demand for NAD, which is the donor substrate for ADP-ribosylation, due to increased PARP activity in response to DNA damage (Pascal, 2018).

## MATERIALS AND METHODS

### Cell culture and stimulation

Y1 cells were grown in DMEM media supplemented with 10% serum, 100 mg/L streptomycin, 25 mg/L ampicillin, 1.2 g/L sodium bicarbonate and kept at 37°C in a 5% CO_2_ incubator. After plating, cells were allowed to reach 30% confluence before serum-starvation for 48 hours. Starved cells were then stimulated with fetal bovine serum (10%) or serum plus FGF-2 (10 ng/mL) and harvested at the indicated timepoints.

### Proteomics and phospho-proteomics

Y1 cells were lysed in a buffer containing 6 M urea, 2 M thiourea, and 60 mM ammonium bicarbonate, pH 8.2, supplemented with phosphatase and protease inhibitors. Proteins were reduced using 10 mM DTT for 1 h at room temperature, alkylated with 20 mM iodoacetamide (IAA) in the dark for 30 min at room temperature, and then digested with trypsin (enzyme:sample ratio, 1:100) overnight at room temperature. Phosphopeptide enrichment was performed using titanium dioxide (TiO2) chromatographic resin, as previously described (Thingholm & Larsen, 2016). Samples were desalted and resuspended in 0.1% formic acid for analysis (2 μL injections) by nLC-MS/MS, composed of a Dionex nano-LC (Thermo Scientific) coupled to a QE-HF Orbitrap mass spectrometer (Thermo Scientific) operating in data-dependent acquisition mode. Chromatography was performed at a flow rate of 300 nL/min over nano-columns (75 μm ID x 25 cm) packed with Reprosil-Pur C_18_-AQ (3 μm, Dr. Maisch GmbH). Water and 80% acetonitrile, both containing 0.1% formic acid, served as solvents A and B, respectively. The gradient for proteome analysis consisted of 5-36% solvent B for 71 min, 36-65% B for 5 min, 65-95% B for 0.1 min, and 10 min of isocratic flow at 95% B. The gradient for phospho-proteome analysis consisted of 2-28% B for 61 min, 28-38% B for 19 min, 38-85% B for 1 min, and 15 min of isocratic flow at 85% B. A full MS scan was acquired over 300-1200 *m/z* in the Orbitrap in centroid mode with a resolution of 60K, AGC target of 1e6, and maximum injection time of 100 ms. The top 20 most intense ions (charge state 2+ and higher) were selected for MS/MS by high-energy collision dissociation (HCD) at 27 NCE, and fragmentation spectra were acquired in the Orbitrap in centroid mode with a resolution of 15K, AGC target of 5e4, and maximum injection time of 150 ms. The proteome analysis used similar settings except the full MS scan was acquired in profile mode and the maximum injection time for MS and MS/MS scans was 50 ms. Data was processed in Proteome Discoverer 2.2 (Thermo Scientific) using the Sequest-HT node to search MS/MS spectra against the UniProtKB/SwissProt (organism: *Mus musculus* / November 2017) database along with a contaminant database. Trypsin was selected as the protease with a maximum of 2 missed cleavages. For the proteome data, the search parameters were as follows: 20 ppm precursor ion mass tolerance; 0.05 Da fragment ion mass tolerance; carbamidomethylation (+57.021 Da to Cys) as a static modification; and oxidation (+15.995 Da to Met) and acetylation (+42.011 Da to protein N-terminus) as variable modifications. For the phospho-proteome data, the search parameters were as follows: 10 ppm precursor ion mass tolerance; 0.02 Da fragment ion mass tolerance; carbamidomethylation (+57.021 Da to Cys) as a static modification; and phosphorylation (+79.966 Da to Ser/Thr/Tyr) as a variable modification. The Percolator node was used with default parameters and data were filtered for < 1% FDR at the peptide level. Statistical analysis was performed in R with the MSstats package (MacLean *et al*, 2014). Pathway analysis was performed using the ReactomePA (Yu & He, 2016) and ClusterProfiler (Yu *et al*, 2012) packages. The experiments were performed with 3 biological replicates.

### Histone extraction, derivatization, and analysis by LC-MS/MS

Histones were extracted and purified from nuclei as described in Sidoli et al. (Sidoli *et al*, 2016). Briefly, nuclei were extracted from Y1 cells (10^7 cells) using nuclear isolation buffer (15 mM Tris, 60 mM KCl, 15 mM NaCl, 5 mM MgCl_2_, 1 mM CaCl_2_, pH 7.5), supplemented with 0.3% NP-40, 1 mM dithiothreitol, 500 μM AEBSF, 5 nM microcystin, and 10 mM sodium butyrate (protease, phosphatase and deacetylase inhibitors). Histones were isolated from nuclei using 0.2 M H_2_SO_4_ and precipitated by the addition of TCA to 33% with incubation overnight on ice. Quantification was performed using the BCA Protein Quantification Assay Kit (Thermo Scientific). Derivatization and digestion of histones and analysis of the resulting peptides were performed as described in Sidoli et al (Sidoli *et al*, 2016). In short, 20 μg of purified histones were resuspended in 100 mM NH_4_HCO_3_. Derivatization reagent was prepared by mixing propionic anhydride with propanol in a 1:3 (v/v) ratio and mixed with histones in a 1:4 (v/v) ratio. The propionylation reaction was performed twice with drying by speed-vac in between. Then, propionylated histones were digested overnight with trypsin (enzyme:sample ratio, 1:20) followed by a second round of two propionylation reactions, as described above. Samples were desalted prior to nLC-MS/MS analysis. Samples were resuspended in 0.1% trifluoroacetic acid (TFA) and injected (1 μL) onto a 75 μm ID x 25 cm nano-column packed in-house with Reprosil-Pur C18-AQ (3 μm, Dr. Maisch GmbH) and attached to an EASY-nLC system (Thermo Scientific). The nano-LC delivered a flow rate of 300 nL/min with a programmed gradient from 5% to 28% B (solvent A = 0.1% formic acid in water; solvent B = 0.1% formic acid in 80% acetonitrile) over 45 minutes, followed by a gradient from 28% to 80% solvent B in 5 minutes and then 10 min of isocratic flow at 80% B. The instrument was coupled in line with an Orbitrap Elite (Thermo Scientific) mass spectrometer running in data-independent acquisition (DIA) mode as previously described (Sidoli *et al*, 2015). The DIA method consisted of one full MS scan (300–1100 *m/z*) in the Orbitrap at a resolution of 120K, followed by 16 DIA MS/MS scans across this mass range in windows of 50 *m/z* with fragmentation by CID (35 NCE) and detection in the ion trap. The DIA scans were punctuated with a second full MS scan after the eighth MS/MS scan. DIA data was analyzed using EpiProfile (Yuan *et al*, 2015). The relative abundances of different modifications on a peptide with a given amino acid sequence were calculated based on the AUC of extracted ion chromatograms of all forms, modified and unmodified, of that peptide. For isobaric peptides, the relative ratio of two isobaric forms was estimated by averaging the ratio for each diagnostic fragment ion having a differential mass between the two species. Statistical analysis was performed in R. The experiment was performed with 4 biological replicates.

### Chromatin Immunoprecipitation Sequencing (ChIP-seq)

Approximately 10^6 cells were crosslinked in 1% paraformaldehyde for 5 mins followed by quenching in 125 mM glycine for 5 mins. The cross-linked cells were lysed as previously described (Lee *et al*, 2006) except that N-lauroylsarcosine was used at 0.1%. Lysates were sonicated with a S220 Focused-ultrasonicator (Covaris) for 15 min. Sonicated lysates were supplemented with 1% Triton X-100 and then cleared by centrifugation at full speed in a microcentrifuge for 10 min at 4°C. The supernatant was transferred to a new tube and the protein content was measured by Bradford assay. Extracts were incubated overnight at 4°C with 5 μg of anti-H3K27ac (Active Motif) prebound to protein G Dynabeads (Thermo Scientific) that were previously blocked in PBS with 0.5% BSA. Beads were washed three times with RIPA buffer (50 mM HEPES-KOH pH 7.5, 500 mM LiCl, 1 mM EDTA, 1% NP-40, 0.7% sodium deoxycholate) and then once with TE buffer containing 50 mM NaCl. Beads were then incubated in elution buffer (50 mM Tris-HCl, pH 8.0, 10 mM EDTA, 1% SDS) for 30 min at 65°C. Crosslink reversal was performed by overnight incubation at 65°C. Samples were diluted with 1 volume of TE buffer and then treated with RNaseA (0.2 mg/ml for 2 hrs at 37°C) and proteinase K (0.2 mg/ml for 2 hrs at 55°C) prior to DNA extraction with phenol:chloroform:isoamyl alcohol. Library preparation was performed using the NEBNext®Ultra™ II DNA Library Prep Kit for Illumina (New England Biolabs), and library quantification was performed using the KAPA library quantification kit (Kapa Biosystems). Paired-end sequencing (75 cycles with a 6-cycle index read) was performed with the NextSeq 500 (Illumina) platform. Sequencing data was aligned to the mm10 genome using STAR 2.5.2a (Dobin *et al*, 2013) (outFilterMultimapNmax 20 --outFilterMismatchNmax 999 --alignMatesGapMax 1000000) and filtered for uniquely aligning reads (MAPQ = 255). Read pairs were collapsed into single fragments and then filtered for a maximum length of 1000 bp. Peak calling was performed using MACS2 (v2.2.6) (Zhang *et al*, 2008) (-f BEDPE -g mm -q 1e-2) for each enrichment with the corresponding input as background. Quantitative comparisons were performed using the DiffBind package (Stark & Brown, 2011) in R. Genome tracks were normalized to input. The experiment was performed for 2 biological replicates.

### RNA sequencing (RNA-seq)

RNA was extracted from cells using the RNeasy mini kit (Qiagen). mRNA isolation, cDNA synthesis, and library preparation were performed using the NEBNext Ultra Directional RNA Library Prep Kit for Illumina (New England Biolabs) according to the manufacturer’s instructions. Library quantification and Illumina sequencing were performed as above. Paired reads were aligned to the mm10 genome using STAR (Dobin *et al*, 2013) (--outFilterType BySJout --outFilterMultimapNmax 20 --alignSJoverhangMin 8 --alignSJDBoverhangMin 1 --outFilterMismatchNmax999 --alignIntronMin 20 --alignIntronMax 1000000 --alignMatesGapMax 1000000) and filtered for unique alignments. Quantification was performed with HTSeq (Pyl *et al*, 2014) (-f sam -r name -s reverse -t exon -i gene_id --nonunique none --secondary-alignments ignore --supplementary-alignments ignore) with subsequent analysis by DESeq2 (Anders & Huber, 2010) in R and the Gene Set Enrichment Analysis (GSEA) tool (Subramanian *et al*, 2005; Mootha *et al*, 2003). For GSEA, genes with low HTSeq counts (< 10 summed across all samples) and undefined adjusted p-values in DESeq2 were omitted. All other genes (11,951) were pre-ranked by log2(FGF/FBS) and analyzed by GSEA. The experiment was performed for 3 biological replicates.

## ACKNOWLEDGEMENTS

The authors thank Ivan Novaski for assistance with cell culture; Dr. Payel Sen, Dr. Marcelo Reis and Dr. Hugo Armelin for revising the manuscript; Greg Donahue and the Berger Lab for assistance with sequencing analysis; the Fundação de Amparo à Pesquisa do Estado de São Paulo (FAPESP) grants #2011/22619-7 (J.P.C.C.), #2018/15553-9 (J.P.C.C.), #2013/07467-1, #2015/04867-4 (M.L.), #2016/24881-4 (M.L.) and #2017/15835-1 (M.L.), and #2017/18344-9 (F.N.L.V.); the Crohn’s and Colitis Foundation (P.J.L.); and the National Institutes of Health (2T32CA009140-41A1 to P.J.L, P01AG031862 to B.A.G., R01CA196539 to B.A.G.).

## CONTRIBUTIONS

P.J.L. and M.L. collected data with assistance from S.S., M.C. and F.N.L.V.; P.J.L., M.L. and S.S., performed data analysis; M.L. and P.J.L. generated the figures, M.L. and P.J.L. conceptualized and wrote the manuscript with revisions from all authors, J.P.C.C. participated in initial conceptualization and supervision; P.J.L., S.S., and B.A.G. provided supervision for mass spectrometry analyses; P.J.L. supervised the sequencing experiments.

---

**Supplemental Figure 1.**
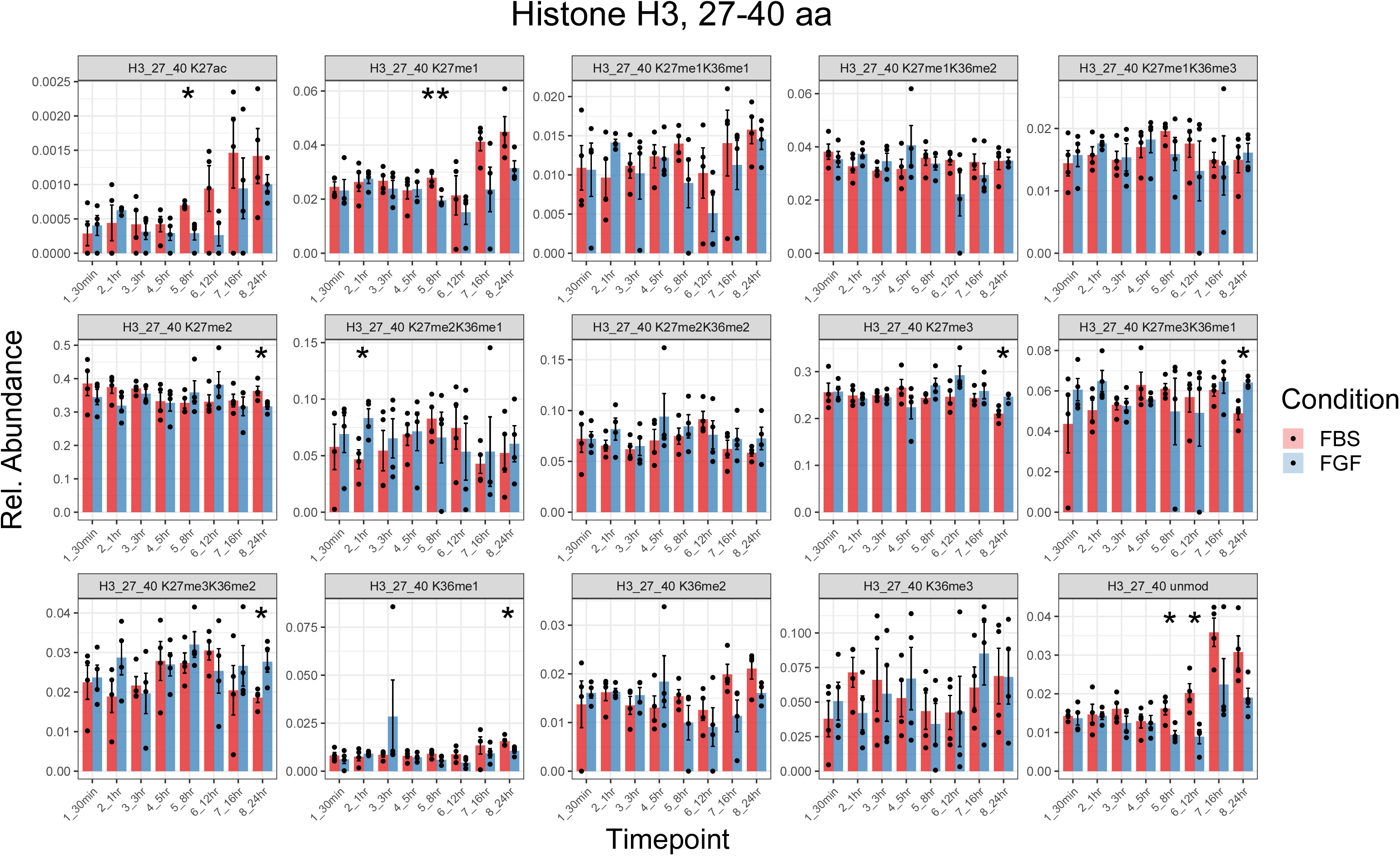
Relative abundances of PTMs on peptide KSAPATGGVKKPHR (aa 27-40) from histone H3. Histones from Y1 cells treated with FBS or FBS + FGF-2 for various amounts of time were digested into peptides and analyzed by LC-MS/MS. The relative abundances of all the uniquely modified forms of the H3 peptide, amino acids 27-40, are plotted over time for both conditions (mean ± sem). * p < 0.05, ** p < 0.01 by unpaired t-test.

**Supplemental Figure 2.**
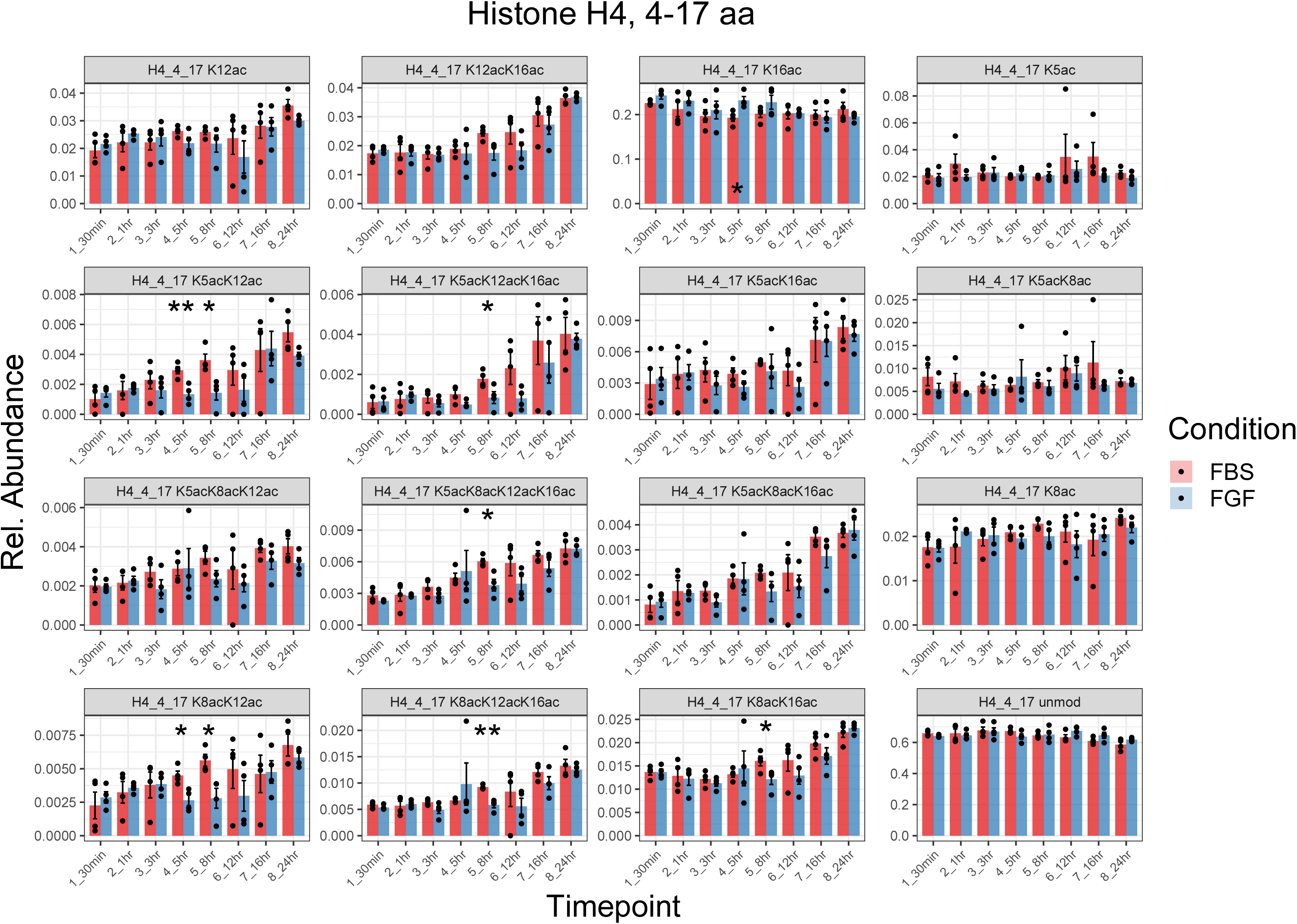
Relative abundances of PTMs on peptide GKGGKGLGKGGAKR (aa 4-17) from histone H4. Histones from Y1 cells treated with FBS or FBS + FGF-2 for various amounts of time were digested into peptides and analyzed by LC-MS/MS. The relative abundances of all the uniquely modified forms of the H4 peptide, amino acids 4-17, are plotted over time for both conditions (mean ± sem). * p < 0.05, ** p < 0.01 by unpaired t-test.

**Supplemental Figure 3.**
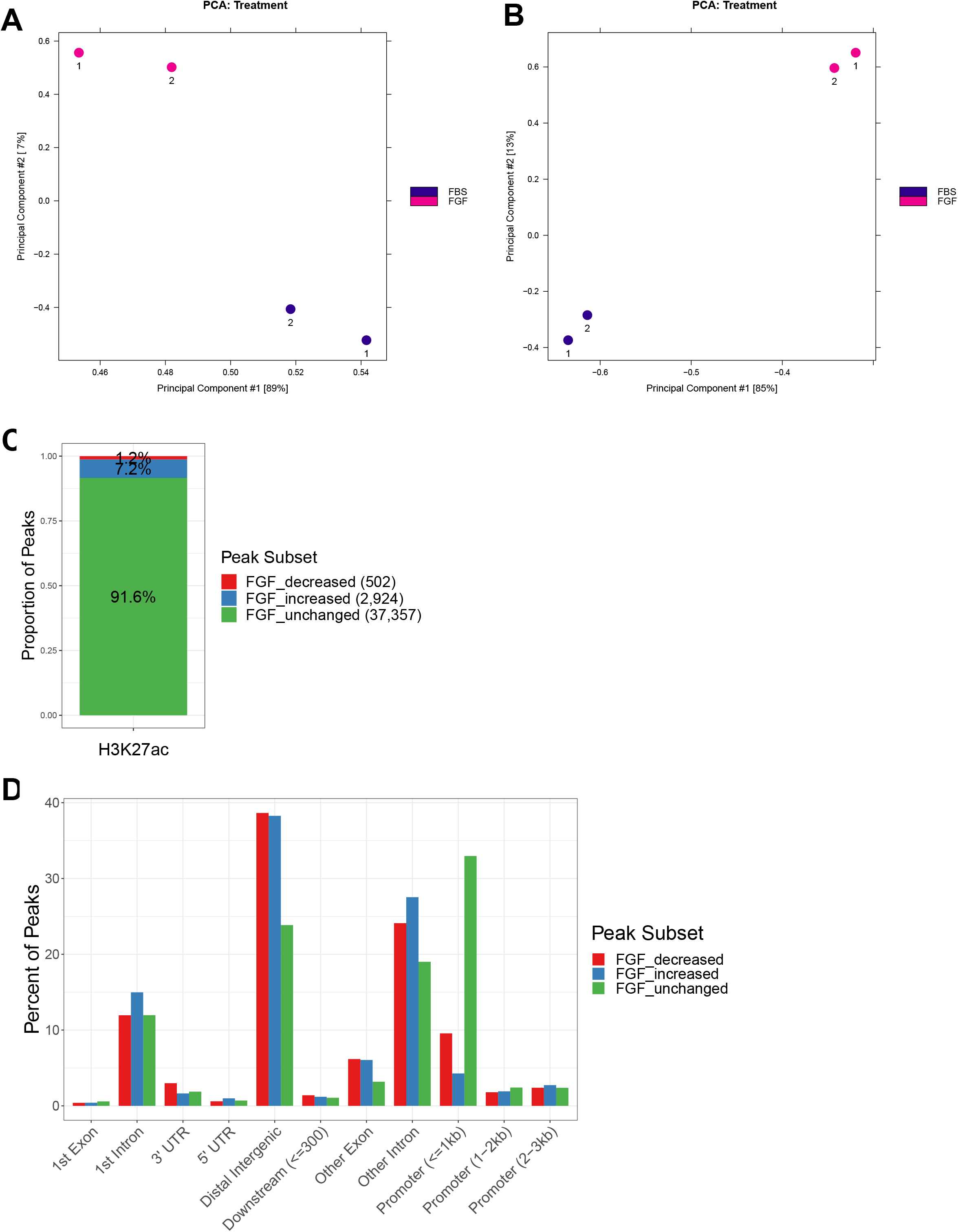
Summary of H3K27ac ChIP peaks and their genomic locations. A) After alignment and processing, H3K27ac ChIP-seq data was analyzed in R with the DiffBind (Stark & Brown, 2011), ChIPseeker (Yu *et al*, 2015), and ClusterProfiler (Yu *et al*, 2012) packages. Principal component analysis (PCA) was used to cluster the samples based on 40,783 consensus H3K27ac peaks. Each point represents an individual replicate, labeled as 1 or 2, from the FBS or FGF condition after 5 hrs of treatment. B) PCA was also performed considering only the 3,246 differential H3K27ac peaks (q < 0.05, absolute log2 fold-change > 1). C) H3K27ac peaks were classified as having unchanged, increased (q < 0.05, log2 fold-change > 1), or decreased (q < 0.05, log2 fold-change < -1) intensity in the FGF condition compared to the FBS control after 5 hrs. The proportion of peaks in each subset is plotted. Numbers in parentheses indicate the total number of peaks in each category. D) H3K27ac peaks were further classified based on their proximity to annotated genomic features. The percent of H3K27ac peaks from each subset is presented according to their genomic annotation.

**Supplemental Figure 4.**
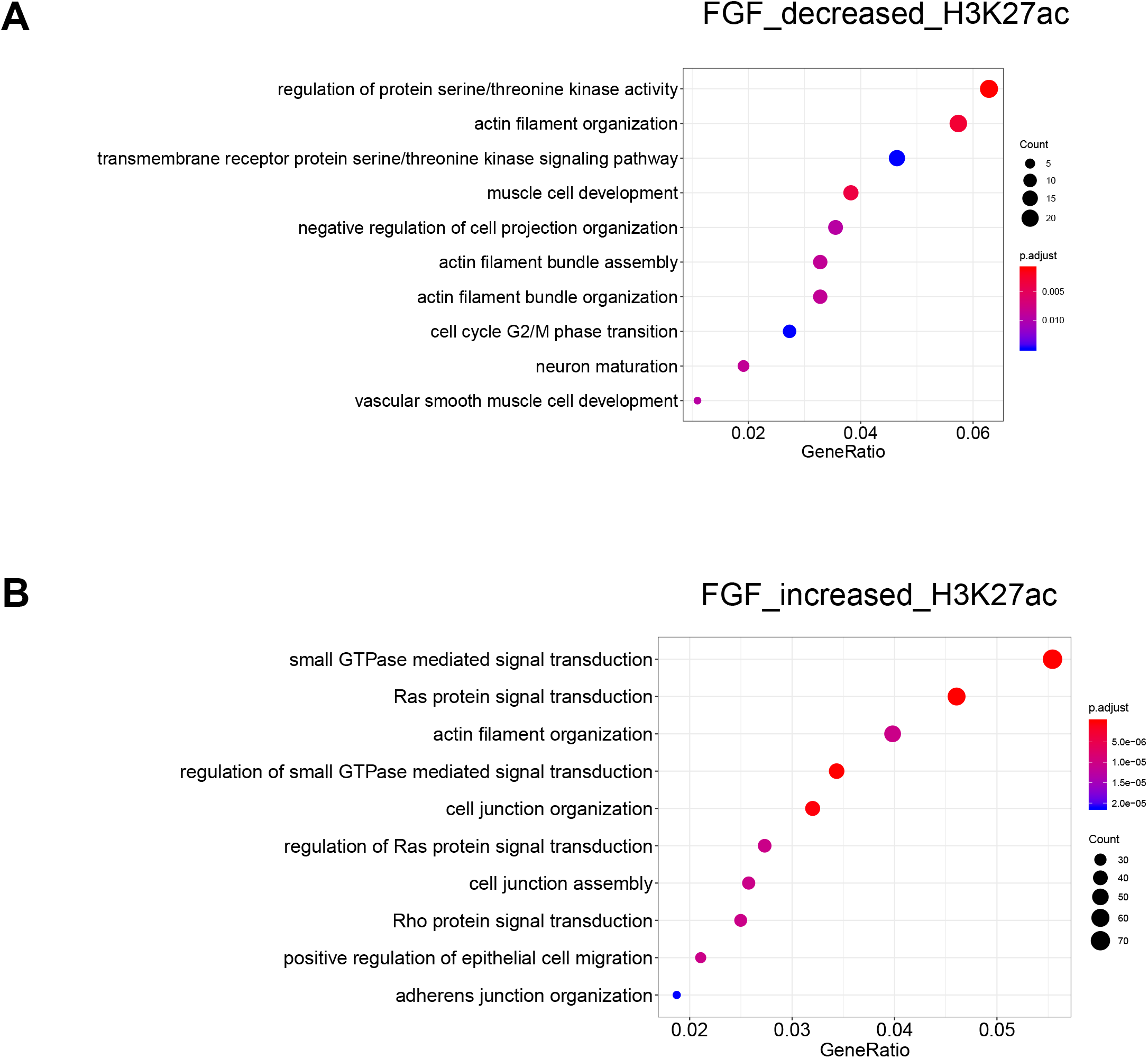
Gene ontology analysis of H3K27ac peaks with differential abundance after FGF-2 treatment. A,B) H3K27ac peaks with quantitative differences between the FGF treatment and the FBS control were annotated with the nearest gene. GO enrichment analysis was performed for peaks with lower intensity in the FGF condition (A, 502 peaks representing 415 genes, log2(FGF/FBS) < -1 at FDR < 0.05) and separately for peaks with higher intensity in the FGF condition (B, 2924 peaks representing 1421 genes, log2(FGF/FBS) > 1 at FDR < 0.05). The top 10 GO terms are displayed. The entire genome served as background.

**Supplemental Figure 5.**
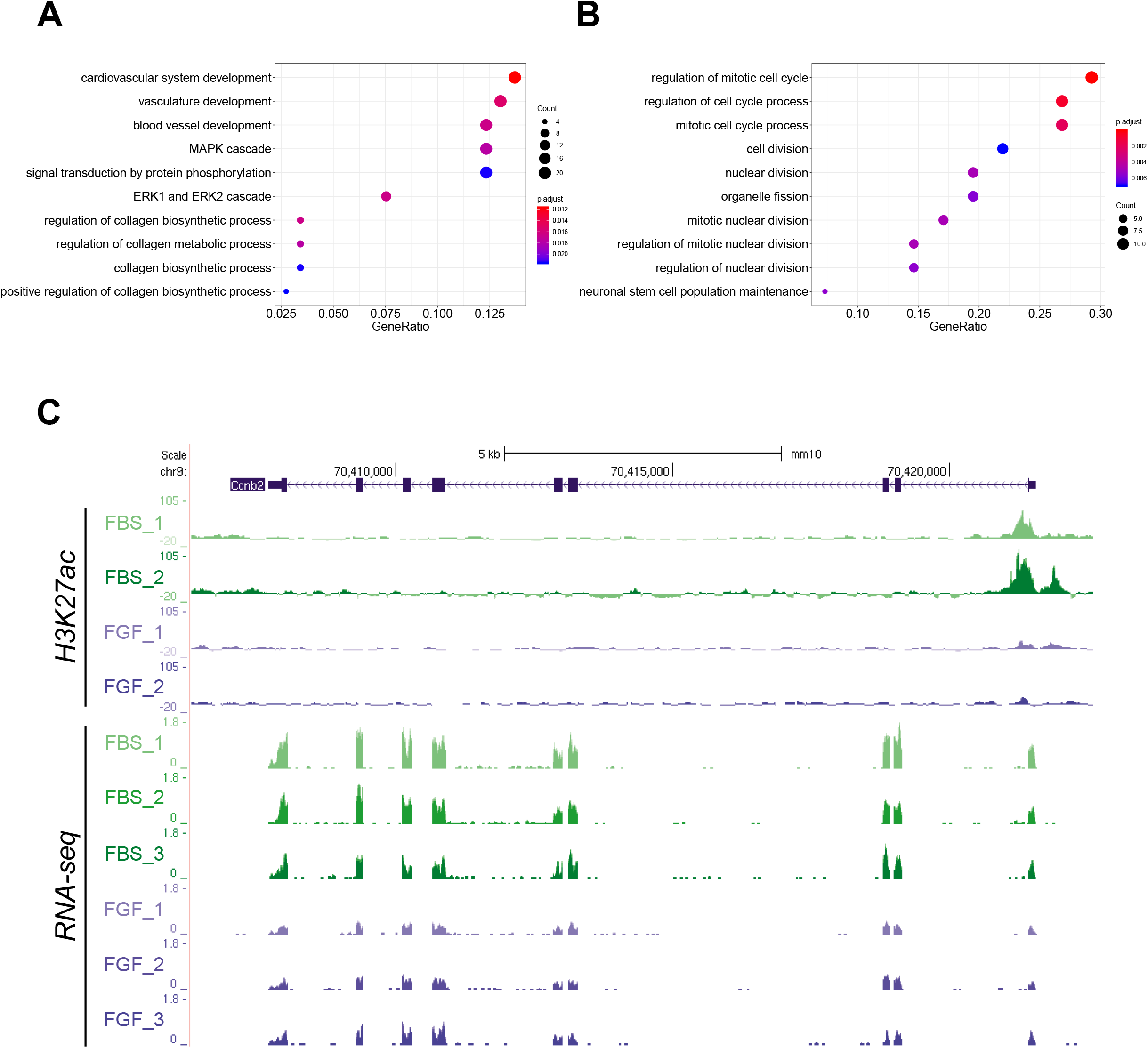
Gene ontology analysis of genes with matching trends in levels of RNA and H3K27ac. A,B) Gene ontology analysis was performed on genes showing concordant trends in RNA and H3K27ac levels. Enriched terms are presented for (A) genes with increased RNA expression and H3K27ac intensity (157 genes with H3K27ac log2(FGF/FBS) > 0 at FDR < 0.05 and RNA log2(FGF/FBS) > 0 at p-adj < 0.05) and for (B) genes with decreased RNA expression and H3K27ac intensity (42 genes with H3K27ac log2(FGF/FBS) < 0 at FDR < 0.05 and RNA log2(FGF/FBS) < 0 at p-adj < 0.05). C) RNA-seq and H3K27ac ChIP-seq tracks from the FBS (green) and FGF (blue) conditions are shown for the genomic region near the *Ccnb2* gene (cyclin-B2).

**Supplemental Figure 6.**
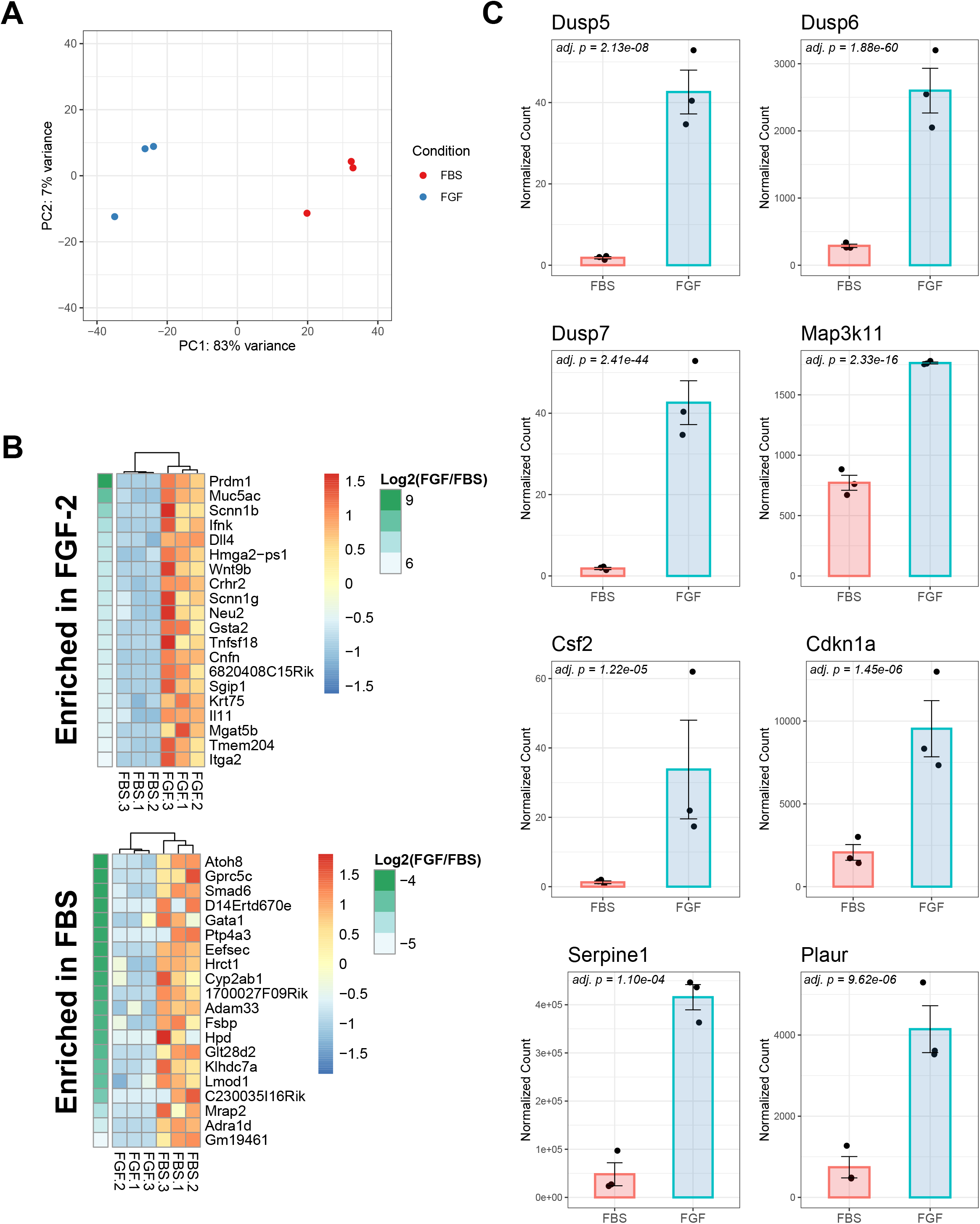
Summary of RNA-seq results and plots of selected genes demonstrating differential expression. A) Replicates from the RNA-seq analysis were clustered by PCA (11,952 genes with valid adjusted p-values). Each point represents an individual replicate. The FBS condition is shown in red while the FGF condition is shown in blue. B) Heatmaps of the 20 genes showing the largest changes in log2(FGF/FBS) (p.adj < 0.05) are presented. C) Normalized RNA-seq counts are presented for several genes related to MAPK signaling, cell cycle regulation, and the senescence-associated secretory phenotype. Bar charts display the group mean ± s.e.m. with individual replicates as points. Adjusted p-values are shown from the DESeq2 analysis.

**Supplemental Figure 7.**
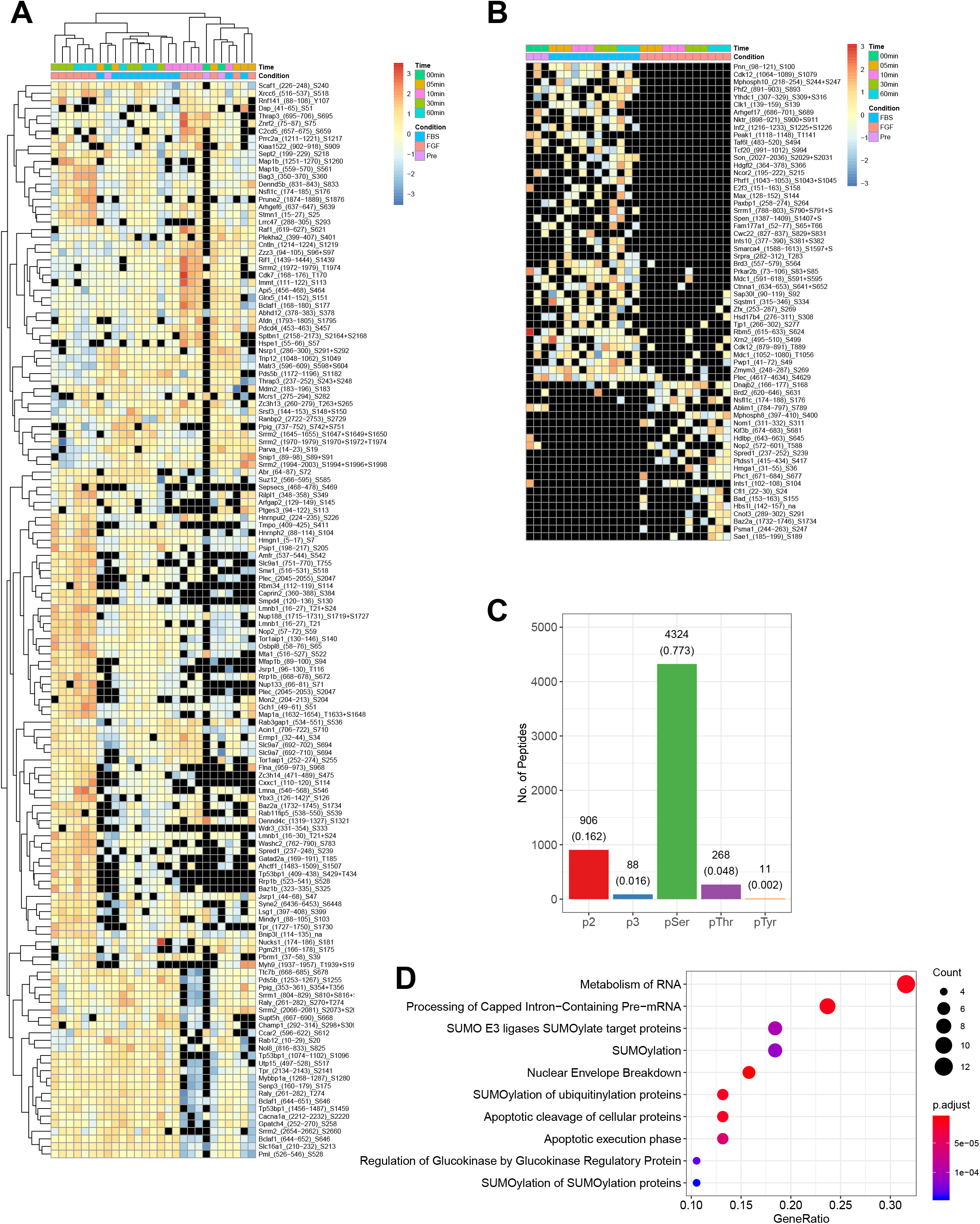
Summary of phospho-proteome results and additional analyses. A) A heatmap of phospho-peptides with significant changes (p < 0.05, abs(Z-score of log2(FGF/FBS)) > 2, quantified in all replicates in both conditions for at least 1 timepoint) in intensities is presented. Unlike Fig. 4A, phospho-peptide intensity is not normalized to the intensity of the respective protein. Missing values are shown in black. Row names indicate the gene symbol, starting and ending positions of the peptide, and the phosphorylation sites. The asterisk for Ybx3 indicates an ambiguous protein assignment due to an identical sequence in Ybx1. The na for Bnip3l indicates an ambiguous phospho-serine assignment. B) A heatmap of phospho-peptides with significant enrichment in either the FBS or FGF condition is presented. Peptides were selected based on statistical significance (p < 0.05) based on a chi square test applied to the number of detections in the FGF and FBS conditions, regardless of timepoint. Missing intensity values are represented in black. Row names indicate the gene symbol, starting and ending positions of the peptide, and the phosphorylation sites. Entries with na or S without a position represent ambiguous phospho-site assignments. C) The numbers of phospho-peptides (total = 5,597) bearing single phosphorylation events at serine, threonine, or tyrosine are plotted as well as those bearing two or three simultaneous phosphorylation events. The frequency is shown in parentheses below the absolute number of unique phospho-peptides. The phospho-peptide HEpSSEEGDSHRR is not included in the counts. D) Pathway analysis (Reactome) was performed on proteins (n = 67) with protein-normalized phospho-peptides showing significant changes across conditions (p < 0.05 and absolute value of Z-score of log2[FGF/FBS] > 2 at 5, 10, 30, or 60 mins).

**Supplemental Figure 8.**
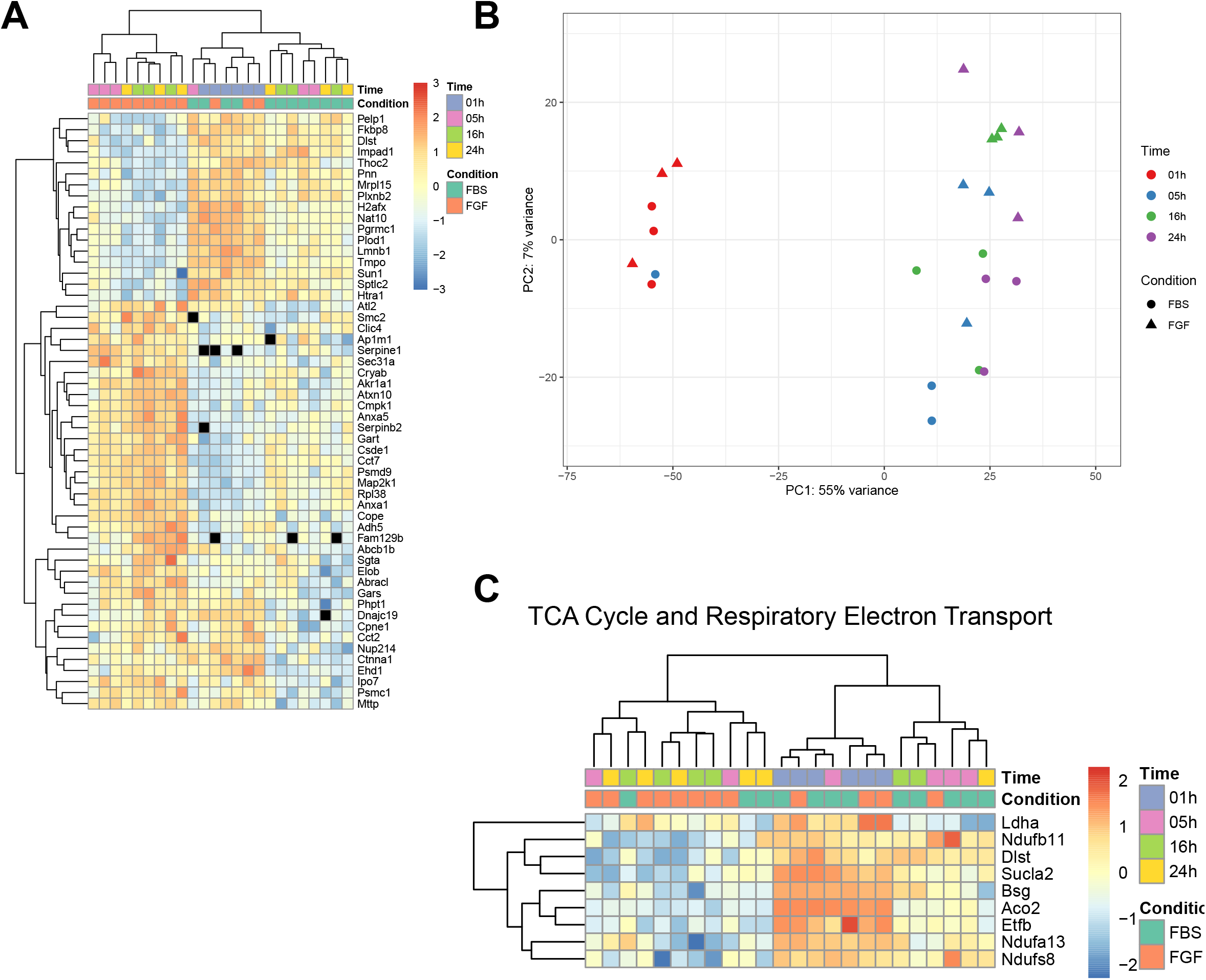
Summary of proteome results. A) A heatmap is presented showing proteins with significant changes in intensity across conditions (p < 0.05 for both 16 and 24 hrs with consistent sign in Z-scores of log2[FGF/FBS] at 5, 16, and 24 hrs). Missing values are shown in black. B) Proteins with complete sets of measurements across all conditions and replicates (n = 1742) were used for PCA. Individual replicates are plotted with a different color representing each timepoint and a different shape representing each condition. C) Proteins related to the TCA cycle by GO analysis and showing significant changes (p < 0.05 at 16 or 24 hrs for log2[FGF/FBS] and consistent sign in Z-score of log2[FGF/FBS] at 5, 16, and 24 hrs) across conditions are presented in a heatmap.

**Supplemental Figure 9.**
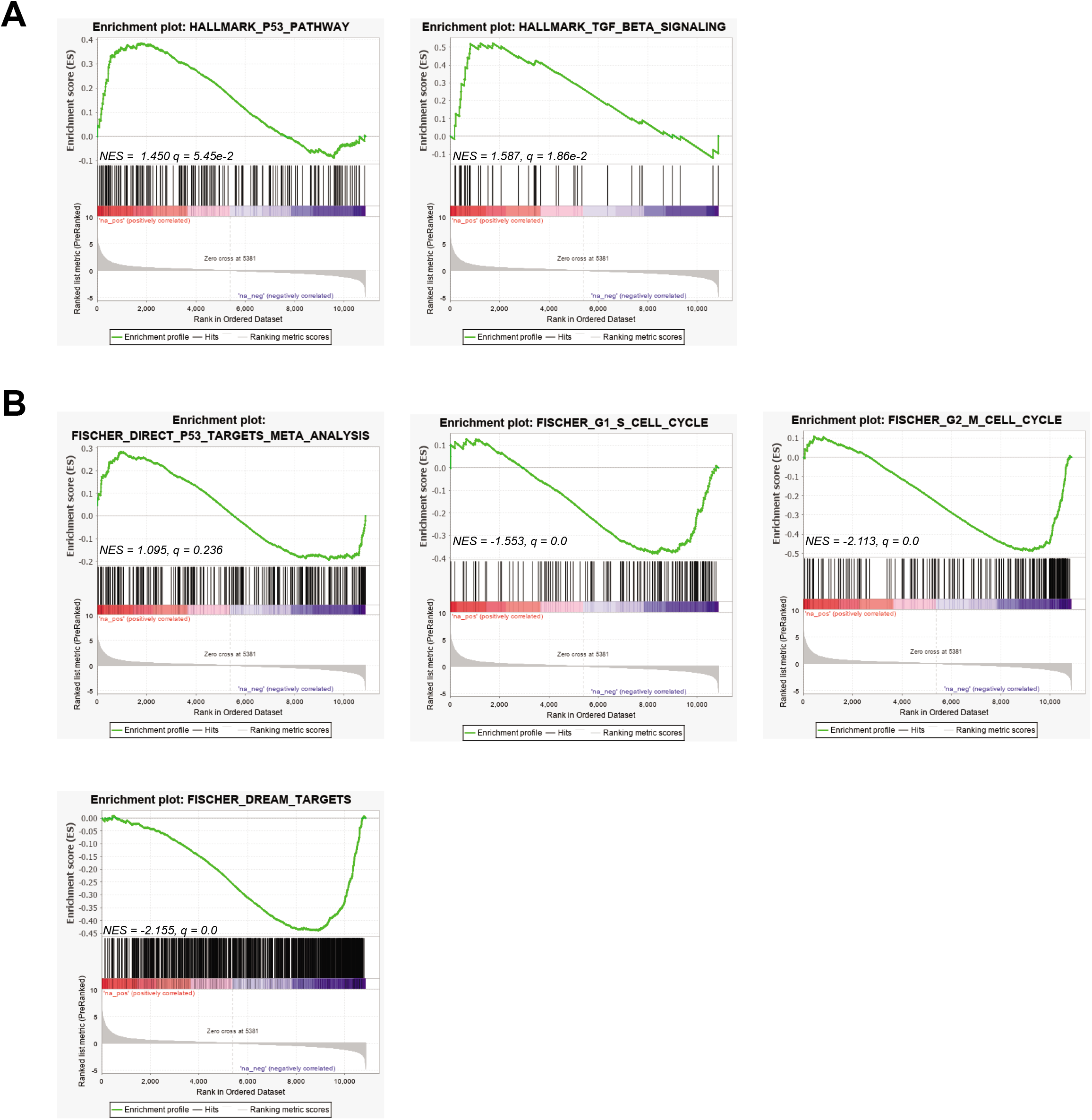
GSEA of p53, TGFb, DREAM, and cell cycle pathways. A) Enrichment plots for the hallmark p53 and TGFb pathways from the initial GSEA. B) Enrichment plots from a targeted GSEA focusing on direct p53 transcriptional targets, DREAM targets, and genes with peak expression at the G1/S and G2/M transitions based on a published meta-analysis (Fischer *et al*, 2016a).

**Table 1. Analysis of H3K27ac ChIP-seq data**

This table reports qualitative and quantitative information regarding the consensus H3K27ac peaks that were called by MACS2 and processed by DiffBind in R. Genomic annotation was provided by the ChIPseeker package in R. Peaks were assigned to FGF_up, FGF_down, and Unchanged subsets based on FDR < 0.05 and Fold (calculated as log2[FGF/FBS]) > 1 or < -1. The Unchanged subset contains 37,346 peaks instead of 37,357 due to 11 peaks with ChrUn designation not receiving a genomic annotation. FBS1 and FBS2 represent the two replicates from the FBS condition while FGF1 and FGF2 represent the two replicates from the FGF-2 condition.

**Table 2. Integration of RNA-seq with H3K27ac ChIP-seq.**

This table reports information regarding the integrated analysis of the H3K27ac ChIP-seq and RNA-seq datasets after separate processing by DiffBind and DESeq2. H3K27ac peaks were required to be within 10 kb of the TSS. For genes with multiple H3K27ac peaks, the peak closest to the TSS was selected and in the case of ties, the most intense peak was selected. Genes were also required to have defined adjusted p-values and fold-changes from the DESeq2 analysis. Genes with concordant trends (FDR < 0.05 and log2[FGF/FBS] > 0 or < 0 for both RNA-seq and ChIP-seq) were used for GO analyses.

**Table 3. Analysis of RNA-seq data**

This table reports information regarding the analysis of the RNA-seq data performed with DESeq2. The log2 fold-change indicates log2(FGF/FBS).

**Table 4. Intensities of phospho-peptides before and after normalization to protein intensities**

This table reports the run-normalized intensity measurements from MsStats for the identified phospho-peptides, which are uniquely identified based on concatenation of the protein accession, peptide sequence, and modification site assignments (AccessionModSeq). The phospho-peptide values are matched to run-normalized protein intensity measurements across the rows with the final set of columns representing protein-normalized phospho-peptide intensities. The peptides have not been filtered for consistent detection across replicates. Additional peptide information (ambiguity, accessions, modifications, missed cleavages, sequence) originated from ProteomeDiscoverer. The peptide HEpSSEEGDSHRR has no intensity measurements after processing by MsStats, most likely due to low intensities reported by ProteomeDiscoverer.

**Table 5. Statistical analysis of phospho-peptides with normalization to protein levels**

This table reports statistics, based on MsStats and other tools in R, for the identified phospho-peptides, which have been normalized to the intensity of their respective protein and are uniquely identified based on concatenation of the protein accession, peptide sequence, and modification site assignments (AccessionModSeq). Significance values are derived from unpaired Student’s t-tests on intensity values from MsStats. N indicates the number of replicates in which the phospho-peptide was detected. Additional peptide information (ambiguity, accessions, modifications, missed cleavages, sequence) originated from ProteomeDiscoverer.

**Table 6. Statistical analysis of phospho-peptides without normalization to protein levels**

This table reports statistics, based on MsStats and other tools in R, for the identified phospho-peptides, which have not been normalized to the intensity of their respective protein and are uniquely identified based on concatenation of the protein accession, peptide sequence, and modification site assignments (AccessionModSeq). Significance values are derived from unpaired Student’s t-tests on intensity values from MsStats. N indicates the number of replicates in which the phospho-peptide was detected. Additional peptide information (ambiguity, accessions, modifications, missed cleavages, sequence) originated from ProteomeDiscoverer.

**Table 7. Statistical analysis of phospho-peptides based on number of detections across timepoints and conditions**

This table reports information regarding the chi square analysis of phospho-peptides. For each peptide, a contingency table was constructed to test for skewed partitioning of the number of detections in replicates of a condition independent of time (FBS or FGF; 12 detections possible for each) or of a timepoint independent of condition (5 min, 10 min, 30 min, or 60 min; 6 detections possible for each).

**Table 8. Normalized protein intensities from proteome analysis**

This table reports information regarding normalized protein intensity values as calculated by MsStats.

**Table 9. Statistical analysis of proteome data**

This table reports information regarding the statistical analysis of the proteome data performed with MsStats and additional tools in R. NA values originate from proteins with undefined foldchanges at a given timepoint.

